# The transcription regulator Lmo3 is required for cell fate specification in the external globus pallidus

**DOI:** 10.1101/2022.05.24.493171

**Authors:** Shiona Biswas, C. Savio Chan, John L.R. Rubenstein, Lin Gan

**Affiliations:** The Neuroscience Graduate Program, University of Rochester School of Medicine and Dentistry, Rochester, NY 14627; Department of Neuroscience, University of Rochester School of Medicine and Dentistry, Rochester, NY 14627; Department of Ophthalmology and the Flaum Eye Institute, University of Rochester School of Medicine and Dentistry, Rochester, NY 14627; Department of Neuroscience, Feinberg School of Medicine, Northwestern University, Chicago, Illinois, 60611; Department of Psychiatry, UCSF Weill Institute for Neurosciences, University of California at San Francisco, CA 94158; Department of Neuroscience and Regenerative Medicine, Augusta University Medical College of Georgia, Augusta, GA 30912.

## Abstract

The external globus pallidus (GPe) is an essential component of the basal ganglia, a group of subcortical nuclei that are involved in control of action. Changes in the firing of GPe neurons are associated with both passive and active body movements. Aberrant activity of GPe neurons has been linked to motor symptoms of a variety of movement disorders, such as Parkinson’s Disease, Huntington’s disease and dystonia. Recent studies have helped delineate functionally distinct sub types of GABAergic GPe projection neurons. However, little remains known about specific molecular mechanisms underlying the development of GPe neuronal subtypes. We show that the transcriptional regulator Lmo3 is required for the development of medial ganglionic eminence derived Nkx2.1^+^ and PV^+^ GPe neurons, but not FoxP2^+^ neurons or Npas1^+^ neurons. As a consequence of the reduction in PV^+^ neurons, *Lmo3*-null mice have a reduced pallidal input to the subthalamic nucleus.

**SIGNIFICANCE STATEMENT:** The external globus pallidus (GPe) is a critical component of the basal ganglia and can coordinate neuronal activity across the basal ganglia by virtue of its widespread projections to almost all other basal ganglia nuclei. Aberrant activity of GPe neurons has been linked to motor symptoms of a wide variety of movement disorders. Recent advances have delineated functionally distinct sub types of GABAergic GPe projection neurons. However, little remains known about molecular mechanisms underlying their development. Here, we demonstrate that the transcription regulator Lmo3 is required for the development of specific subtypes of GPe neurons, and for their appropriate connectivity with other parts of the basal ganglia.

## Introduction

The external globus pallidus (GPe) is a key component of the basal ganglia. By virtue of its widespread projections to all basal ganglia nuclei (Kita and Kitai, 1994; Bevan, 1998; Kita, 2007; Mallet et al., 2012; Abdi et al., 2015; Saunders et al., 2016; Glajch et al., 2016), as well as other brain regions like the cortex and thalamus (Mastro et al., 2014; Saunders et al., 2015; Abecassis et al., 2020; Courtney et al., 2021; Lilascharoen et al., 2021), it is in a unique position to influence the processing of motor information. Perturbations in the firing pattern and firing rate of GPe neurons have been linked to motor symptoms of a variety of movement disorders, such as Parkinson’s Disease (Pan and Walters, 1988; Nini et al., 1995; Magnin et al., 2000; Magill et al., 2001; Starr et al., 2005; Kita, 2007; Mallet et al., 2008; Chan et al., 2011; Hegeman et al., 2016), Huntington’s disease, and dystonia (Starr et al., 2005, 2008; Nambu et al., 2011; Waldvogel et al., 2014; Hegeman et al., 2016). Work from across several decades had suggested the existence of neuronal heterogeneity within the GPe, with respect to the molecular and morphological type (Iwahori and Mizuno, 1981; Hontanilla et al., 1994; Kita, 1994; Bevan, 1998; Hoover and Marshall, 1999), anatomical features (Iwahori and Mizuno, 1981; Walker et al., 1989; Kita, 1994; Kita and Kitai, 1994; Bevan, 1998; Hoover and Marshall, 1999; Cooper and Stanford, 2000) and electrophysiological characteristics (DeLong, 1971; Walker et al., 1989; Cooper and Stanford, 2000; Mallet et al., 2008). Despite this, progress was slow in understanding the cellular organization and connectivity of this structure, due to the lack of enough objective criteria that could be consistently used to define and distinguish between specific subtypes of GPe neurons (Hegeman et al., 2016).

More recently, studies have made significant progress, demonstrating that the majority of GPe projection neurons, which are GABAergic, can be divided into several different subtypes. Neurons expressing the calcium binding protein parvalbumin (PV) and those expressing the transcription factor Nkx2.1 project primarily to the subthalamic nucleus (STN), and display faster and more regular firing rates *in vivo* and in *ex vivo* slice preparations, while another group of neurons that express the transcription factor FoxP2 and the opioid precursor preproenkephalin, and those expressing the transcription factor Npas1; project heavily to the dorsal striatum and typically display lower and less regular firing rates (Mallet et al., 2012; Mastro et al., 2014; Abdi et al., 2015; Dodson et al., 2015; Hernández et al., 2015; Abrahao and Lovinger, 2018; Abecassis et al., 2020; Ketzef and Silberberg, 2021). The afferent input received by different GPe neurons from other parts of the basal ganglia also differs in type and strength (Pamukcu et al., 2020; Aristieta et al., 2021; Ketzef and Silberberg, 2021; Cui et al., 2021a). As a result, they encode movement and behavior differently via distinctive changes in their firing (Dodson et al., 2015; Mallet et al., 2016) and their optogenetic manipulations alter locomotor activity in different ways (Pamukcu et al., 2020; Aristieta et al., 2021; Cui et al., 2021b). Finally, the firing of different GPe neuronal subtypes is differentially affected by dopamine loss in animal models of Parkinson’s Disease (Mallet et al., 2008, 2012; Abdi et al., 2015).

It is the genetic programs initiated during brain development, that ultimately generate the distinct cell types that have specialized roles in behavior (Kepecs and Fishell, 2014; Kessaris et al., 2014). In humans, haploinsufficiency or mutations of genes that are associated with neurological disorders can often be linked to brain structures whose development has been shown to be critically dependent on the homologs of the same genes, in mice (Sussel et al., 1999; Pohlenz et al., 2002; Breedveld et al., 2002; Leung and Jia, 2016). A few studies have established the spatiotemporal origins of GPe neurons (Marchand and Lajoie, 1986; Nery et al., 2002; Flandin et al., 2010; Nóbrega-Pereira et al., 2010; Dodson et al., 2015). However, specific molecular factors required for the development of GPe neuronal subtypes remain unexplored. The LIM domain containing transcription regulator *Lmo3* is strongly expressed in the developing GPe during embryogenesis (Long et al., 2009; Flandin et al., 2010, 2011), and required for the development of PV^+^ cortical interneurons (Au et al., 2013), which like many GPe neurons, also originate from the medial ganglionic eminence (MGE) (Flandin et al., 2010; Silberberg et al., 2016). In this study, we have addressed the role of Lmo3 in the development of GPe neuronal subtype.

## Results

### *Lmo3* is expressed in the progenitor zones of the developing globus pallidus

Birthdating experiments in rodents have demonstrated that GPe neurons are born between ∼E10.5 and E13.5, with the PV^+^ subset generated primarily from E10.5 to E12.5, while FoxP2 expressing and Npas1 expressing GPe neurons arise mainly between E11.5 and E13.5 (Marchand and Lajoie, 1986; Nóbrega-Pereira et al., 2010; Dodson et al., 2015; Hu et al., 2017). Furthermore, neuronal subtype identity in the GPe is also partly encoded by place of origin. Nkx2.1^+^ and PV^+^ neurons arise mainly from the MGE while approximately half of the Npas1^+^ neurons are born in the MGE progenitor zone and the rest, which also express FoxP2, are born in the lateral ganglionic eminence (LGE) and/or the caudal ganglionic eminence (CGE) (Sussel et al., 1999; Flandin et al., 2010; Nóbrega-Pereira et al., 2010; Dodson et al., 2015; Silberberg et al., 2016). Fate mapping experiments have also revealed the preoptic area (PoA) as the source of a small fraction (∼10%) of GPe neurons, the majority of which express PV and Nkx2.1 (Nóbrega-Pereira et al., 2010; Abecassis et al., 2020).

Thus, we carried out an expression analysis of *Lmo3* via *in situ hybridization* at various embryonic time points and detected *Lmo3* expression from E11.5 onwards throughout the subventricular zone (SVZ) of the MGE, LGE, PoA, and the developing globus pallidus (GPe). (Fig. 1*A*-*D*, *A’*-*C’*, *a’*-*c’*) *Lmo3* expression is maintained in the GPe as well as neighboring regions of the basal ganglia like the striatum, at birth (Fig. 1*E*) and weakly expressed at P30 (Fig. 1*F*, *f*). More recently, single-cell RNA-sequencing experiments have shown that *Lmo3* is expressed in MGE cells differentiating into GPe neurons and cortical interneurons (Su-Feher et al., 2022).

**Figure 1.**
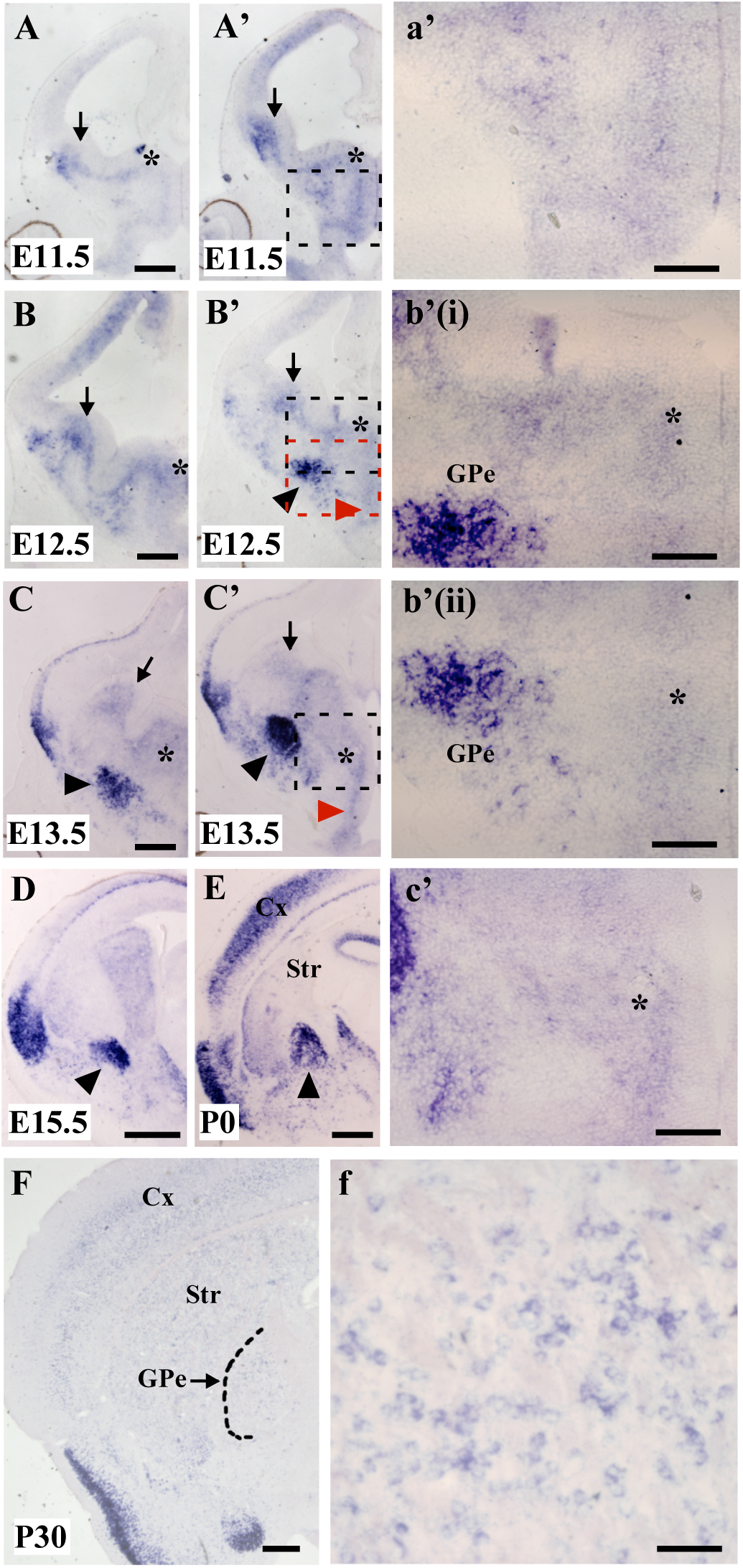
*Lmo3* is expressed in the ganglionic eminences and developing globus pallidus. *Lmo3* is expressed in the SVZ and mantle zone of the LGE (arrows) and MGE (asterisks) in a more rostral (left column) and caudal level (right column), at E11.5 (**A**, **A’**), E12.5 (**B**, **B’**) and E13.5 (**C**, **C’**), in the SVZ of the PoA (red arrowheads) (**B’**, **C’**) and in the developing GPe (black arrowheads) (**B’**, **C**, **C’**). **a’** is a higher magnification view of the boxed region in **A’**. **b’(i)** and **b’(ii)** are higher magnification views of the dorsal MGE (black box) and ventral MGE and PoA (red box) respectively, in **B’**. (**c’**) is a higher magnification view of the SVZ of the boxed region in **C’**. **D**, *Lmo3* expression in the developing GPe (arrowhead) at E15.5. **E**, *Lmo3* expression is maintained in the GPe at birth (P0). **F**, *Lmo3* expression is maintained albeit weakly, in the GPe at P30. **f** is a higher magnification view of the GPe in **F**. Ctx, cortex; GPe, external globus pallidus; Str, striatum; SVZ, subventricular zone. Scale bars: A-C, 500 μm; a’-c’,f, D,E, 200 μm; F, 400 μm.

### *Lmo3*-null mutants display a selective reduction of MGE derived GPe neurons

We immunostained *Lmo3*-null mutant brains for PV and a general neuronal marker NeuN at postnatal day (P)30 (Fig. 2 *A*-*H*). As PV^+^ cell density varies along the medial-lateral axis of the GPe (Hontanilla et al., 1994; Kita, 1994), cell counts were obtained from the entire GPe, in tissue planes at 3 different rostro-caudal levels (Fig. 2*I*). In the *Lmo3*-null, PV^+^ neurons were reduced throughout the rostro-caudal extent of the GPe compared to the wildtype (WT) control (rostral_WT_ = 372.9 ± 6.9, rostral_Lmo3-/-_ = 236.5 ± 20.8, *p* = 0.005; middle_WT_ = 301.3 ± 17, middle_Lmo3-/-_ = 163.6 ± 23.7, *p* = 0.005; caudal_WT_ = 187.8 ± 17.4; caudal_Lmo3-/-_ = 84.4 ± 5.4, *p* = 0.001; two-tailed *t* test, *n*=4 mice for both genotypes; Fig. 2 *J*,*K*,*L*). This corresponds to a PV^+^ reduction in the *Lmo3*-null by 36.5 ± 5.6%, 48.5 ± 8.6% and 53.8 ± 5.3% of the values of control for rostral, middle, and caudal levels, respectively. A striking reduction of PV immunostaining of the neuropil was also observed in the *Lmo3*-null (Fig. 2 *A*,*D*,*G*,*H*). The number of neurons marked by NeuN was also found to be decreased in the *Lmo3*-null (rostral_WT_ = 815.3 ± 60.8, rostral_Lmo3-/-_ = 586.2 ± 41.7, *p* = 0.02; middle_WT_ = 687.8 ± 20.1, middle_Lmo3-/-_ = 556.1 ± 12, *p* = 0.001; caudal_WT_ = 483.4 ± 30.7, caudal_Lmo3-/-_ = 378.5 ± 11.8, *p* = 0.02; two-tailed *t* test, *n*=4 mice for both genotypes). In rostral sections, the reduction of NeuN^+^ neurons in the mutants relative to control appeared higher as compared to the reduction of PV^+^ neurons in the mutants relative to control. However, this difference was not significant (NeuN rostral_WT-Lmo3-/-_ = 229.1 ± 43, PV rostral_WT-Lmo3-/-_ = 136.4 ± 21.8, *p* = 0.12; two-tailed *t* test, *n*=4 mice for both genotypes), and middle and caudal sections also displayed a similar reduction in PV^+^ neurons and NeuN^+^ neurons in mutants relative to control. PV^+^ and NeuN^+^ counts in *Lmo3+/-* mice were not significantly different compared to the WT (For PV, rostral_Lmo3+/-_ = 328.3 ± 21.9, middle_Lmo3+/-_ = 258.5 ± 9.3, caudal_Lmo3+/-_ = 195.5 ± 9.8; rostral *p* = 0.17, middle *p* = 0.11, caudal *p* = 0.71; For NeuN, rostral_Lmo3+/-_ = 834.2 ± 41.3, middle_Lmo3+/-_ = 711.5 ± 31.3, caudal_Lmo3+/-_ = 531.5 ± 24; rostral *p* = 0.82, middle *p* = 0.53, caudal *p* = 0.26, two-tailed *t* test, *n*=3 *Lmo3+/-* mice; Fig. 2 *J*,*K*,*L*).

**Figure 2.**
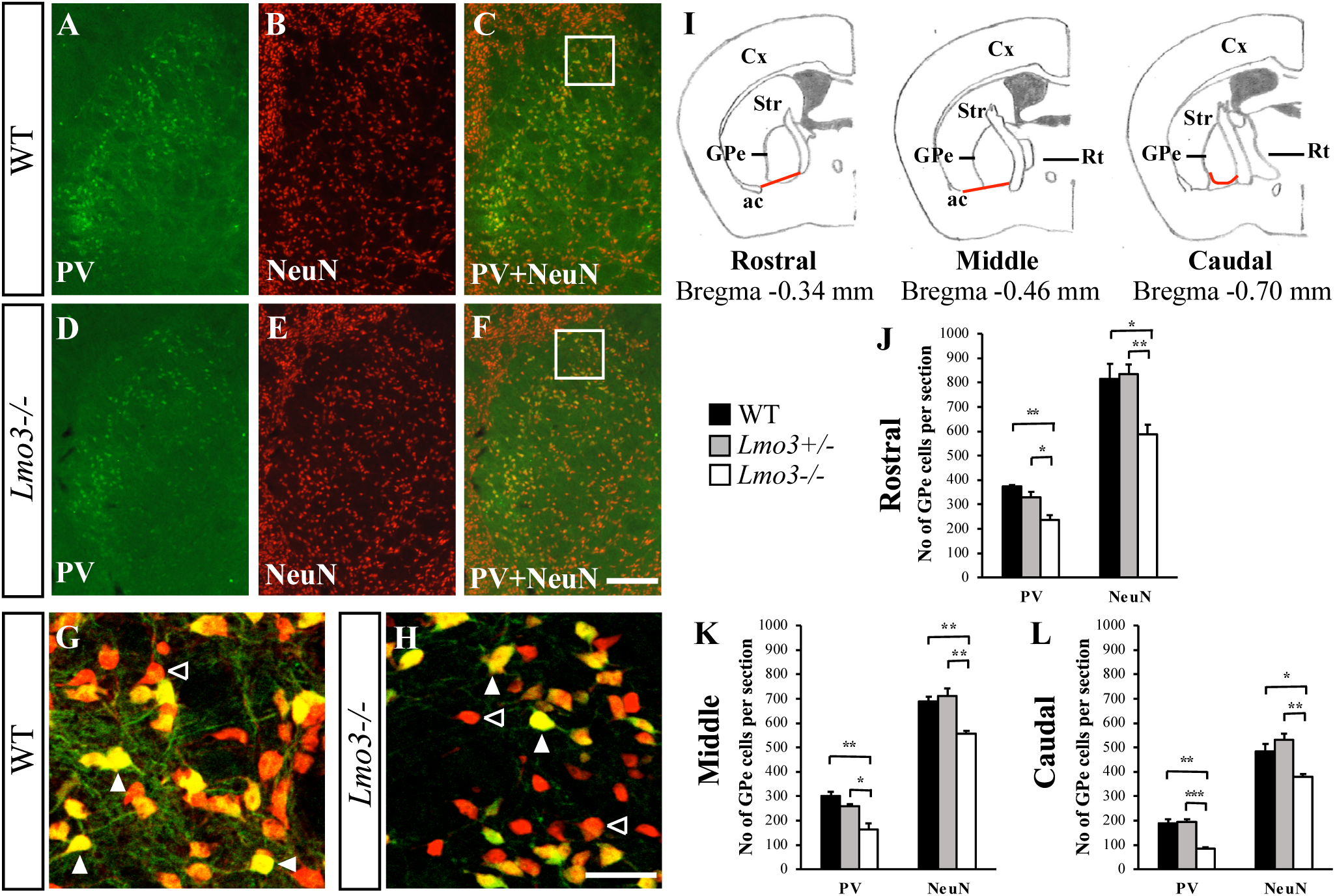
Lmo3 is required for the development of PV-expressing neurons of the external globus pallidus (GPe). **A**-**F**, Coronal views of the GPe of P30 wildtype (**A**-**C**) and *Lmo3*-null mutants (**D**-**F**) immunostained for parvalbumin (PV) and NeuN. **G** and **H** are high magnification images of the regions indicated by the boxes in **C** and **F**, respectively. Filled arrowheads indicate PV^+^;NeuN^+^ neurons; open arrowheads indicate neurons that are only NeuN^+^. **I**, Coronal sections illustrating the rostral, middle and caudal levels of the GPe from which neurons were counted. The ventral GPe borders used for counting were demarcated by the red lines. **J**-**L**, Graphs display the quantification of PV^+^ and NeuN^+^ neurons for the indicated levels. (*n*=4 for WT and *Lmo3-/-* and *n*=3 for Lmo3+/-). [Graphs indicate mean values; error bars show SEM. Two-tailed *t* test, \**p* < 0.05, \*\**p* < 0.01, \*\*\**p* < 0.001] Scale bars: (**F**) 200 μm; (**H**) 50 μm. ac, anterior commissure; Cx, cortex; GPe, external globus pallidus; Rt, reticular nucleus of the thalamus; Str, striatum.

GPe neurons are classified based on their combinatorial expression of several molecular markers: like PV^+^ neurons, all Nkx2.1^+^ neurons originate from the MGE. While most PV^+^ neurons also express Nkx2.1, a subgroup of Nkx2.1^+^ neurons does not express PV (Nóbrega-Pereira et al., 2010; Abdi et al., 2015; Dodson et al., 2015; Abecassis et al., 2020). There can be minor differences between the functional characteristics of a single GPe neuronal subtype depending on the molecular marker it expresses. For example, Nkx2.1^+^;PV^+^ neurons have faster firing rates *in vivo* than Nkx2.1^+^;PV^-^ neurons (Abdi et al., 2015; Dodson et al., 2015). However, both the Nkx2.1^+^;PV^+^ and Nkx2.1^+^;PV^-^ subtypes fire much faster than FoxP2^+^ neurons; and can be distinguished from the latter based on additional electrophysiological properties (Abdi et al., 2015; Dodson et al., 2015; Abrahao and Lovinger, 2018; Abecassis et al., 2020; Aristieta et al., 2021; Ketzef and Silberberg, 2021). To further explore the role of Lmo3 in the development of GPe neuronal subtypes, we performed double immunostaining for Nkx2.1 and FoxP2 at P30 (Fig. 3 *A*-*H*). Nkx2.1^+^ neurons were significantly reduced at all rostro-caudal GPe levels analyzed in the *Lmo3*-null, compared to the WT (rostral_WT_ = 532 ± 56.2, rostral_Lmo3-/-_ = 280.1 ± 25.1, *p* = 0.006; middle_WT_ = 411.3 ± 25.6, middle_Lmo3-/-_ = 215 ± 26.8, *p* = 0.0018; caudal_WT_ = 312.1 ± 14.7, caudal_Lmo3-/-_ = 172.7 ± 18.3, *p* = 0.001; two-tailed *t* test, *n*=4 for both genotypes; Fig. 3 *I*,*J*,*K*). Nkx2.1^+^ neurons in the *Lmo3*-null were reduced by 46.5 ± 5%, 47.8 ± 5.1% and 50.44 ± 4.6% of the values of control for rostral, middle, and caudal levels, respectively. However, the numbers of FoxP2^+^ neurons were unaltered in the *Lmo3*-null, compared to control (rostral_WT_ = 170.6 ± 18.5, rostral_Lmo3-/-_ = 195.3 ± 4.2, *p* = 0.26; middle_WT_ = 158.3 ± 7.8, middle_Lmo3-/-_ = 174 ± 24.4, *p* = 0.57, caudal_WT_ = 142.9 ± 7.4, caudal_Lmo3-/-_ = 182 ± 20.2, *p* = 0.09; two-tailed *t* test, *n*=4 for both genotypes; Fig. 3 *I*,*J*,*K*). Nkx2.1^+^ and FoxP2^+^ counts in *Lmo3+/-* mice were not significantly different compared to the WT (For Nkx2.1, rostral_Lmo3+/-_ = 550 ± 38.3, middle_Lmo3+/-_ = 489.8 ± 53.5, caudal_Lmo3+/-_ = 330.1 ± 9.8; rostral *p* = 0.8, middle *p* = 0.206, caudal *p* = 0.39; For FoxP2, rostral_Lmo3+/-_ = 191.9 ± 4.1, middle_Lmo3+/-_ = 183.8 ± 13.4, caudal_Lmo3+/-_ = 148.4 ± 6.9; rostral *p* = 0.24, middle *p* = 0.18, caudal *p* = 0.62, two-tailed *t* test, *n*=3 *Lmo3+/-* mice; Fig. 2 *J*,*K*,*L*). In agreement with previous studies (Abdi et al., 2015; Dodson et al., 2015) we did not observe any significant co-expression of Nkx2.1 and FoxP2 in the WT, which also held true for the *Lmo3*-null GPe (Fig. 3 *G*,*H*).

**Figure 3.**
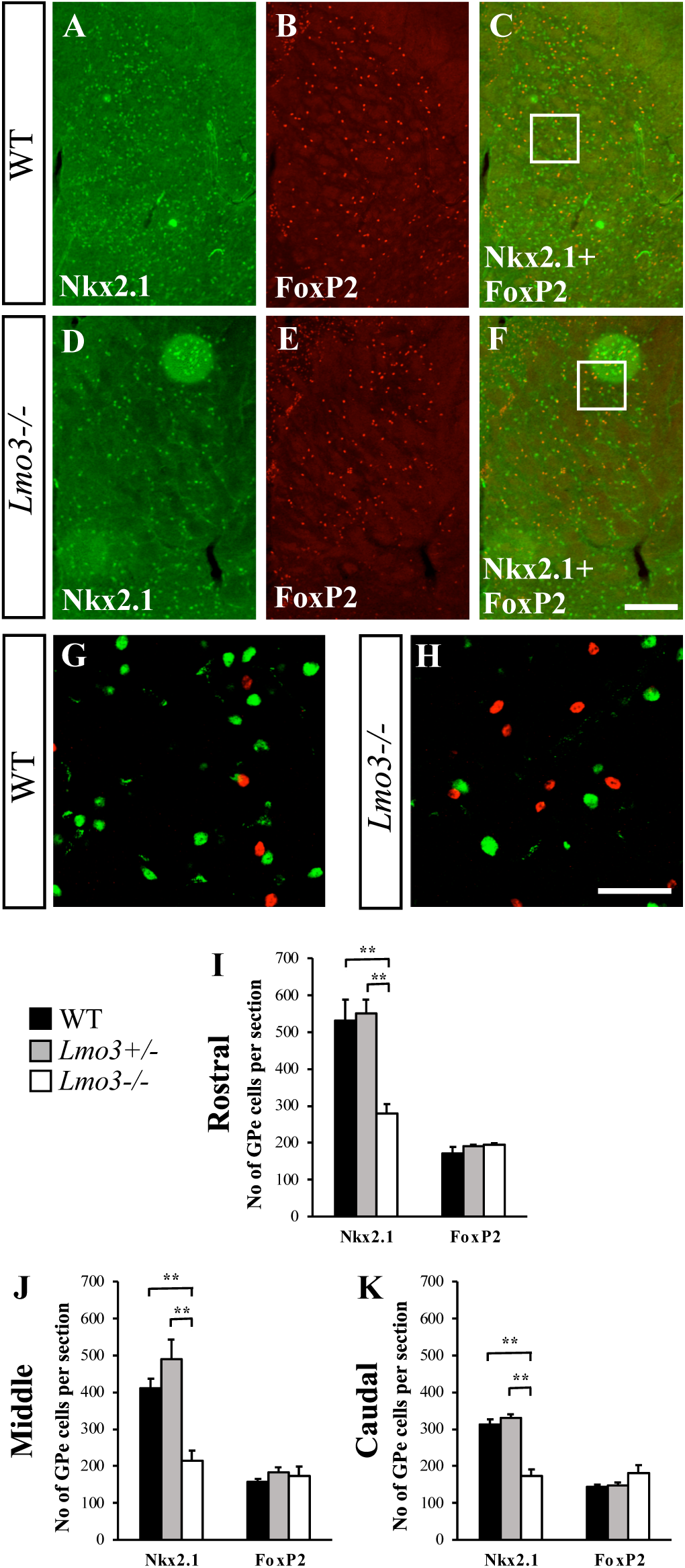
Subtype-specific requirement for Lmo3 in GPe development. **A**-**F**, Coronal views of the GPe of P30 wildtype (**A**-**C**) and *Lmo3*-null mutants (**D**-**F**) immunostained for Nkx2.1 and FoxP2. **G** and **H** are high magnification images of the boxed regions in **C** and **F**, respectively. FoxP2 and Nkx2.1 expression is mutually exclusive in both WT and *Lmo3*-null. **I**-**K**, Quantification of Nkx2.1^+^ and FoxP2^+^ GPe neurons from three different rostrocaudal levels (see Fig. 2. *n*=4 each for WT, *Lmo3+/-* and *Lmo3-/-*). [Graphs indicate mean values; error bars show SEM. Two-tailed *t* test, \**p* < 0.05, \*\**p* < 0.01, \*\*\**p* < 0.001] Scale bars: (**F**) 200 μm; (**H**) 50 μm.

Analysis of Npas1^+^ GPe neurons has shown that they have properties similar to FoxP2^+^ neurons: they send dense axonal projections to the striatum, have significantly lower firing rates compared to PV^+^ neurons, and their optogenetic excitation suppresses locomotion (Hernández et al., 2015; Glajch et al., 2016; Abrahao and Lovinger, 2018; Pamukcu et al., 2020; Cui et al., 2021b). However, Npas1^+^ GPe neurons fall into two categories: those that express FoxP2 and those that do not. Npas1^+^;Foxp2^-^ neurons express Nkx2.1 (these are Nkx2.1^+^;PV^-^ neurons); they originate from the MGE rather than the LGE/CGE (Nóbrega-Pereira et al., 2010; Dodson et al., 2015), and can be distinguished from Npas1^+^;Foxp2^+^ neurons at postnatal ages based on the projection of the former to the cortex (with collaterals in the dorsal striatum) and thalamic reticular nucleus (Abecassis et al., 2020; Cui et al., 2021b). In addition single-cell transcriptomics analysis confirms the existence of separate Npas1^+^;Nkx2.1^+^ and Npas1^+^;Foxp2^+^ neuron clusters within the GPe (Saunders et al., 2018; Cui et al., 2021b). We obtained cell counts from ∼ P30 sections double immunostained for Npas1 and Nkx2.1 (Fig. 4 *A*-*H*); excluding heterozygous mice from further analysis as significant changes in GPe development were not detected in this group. There was no significant change in the number of Npas1^+^ neurons in the *Lmo3*-null (rostral_WT_ = 223.4 ± 6, rostral_Lmo3-/-_ = 238.1 ± 16.1, *p* = 0.418; middle_WT_ = 198.5 ± 13.2, middle_Lmo3-/-_ = 216.2 ± 16.1, *p* = 0.237; caudal_WT_ = 193.7 ± 5.4, caudal_Lmo3-/-_ = 193.9 ± 10.9, *p* = 0.988; two-tailed *t* test, *n*=5 for both genotypes; Fig. 4 *I*). However, we observed a small (∼30%) but significant reduction in the number of Npas1^+^;Nkx2.1^+^ neurons (rostral_WT_ = 67.7 ± 1.7, rostral_Lmo3-/-_ = 46.8 ± 4.3, *p* = 0.011; middle_WT_ = 62.8 ± 3.1, middle_Lmo3-/-_ = 49.2 ± 4.1, *p* = 0.04; caudal_WT_ = 49.9 ± 1.9, caudal_Lmo3-/-_ = 34.6 ± 3.8, *p* = 0.012; two-tailed *t* test, *n*=5 for WT and 4 for *Lmo3*-null for rostral and caudal and *n*=4 for both genotypes for middle; Fig. 4 *J*) which was reflected in a lower percentage of Npas1^+^ neurons co-expressing Nkx2.1^+^ (rostral_WT_ = 30.4 ± 0.88 %, rostral_Lmo3-/-_ = 20.9 ± 1.4 %, *p* = 0.0005; middle_WT_ = 32.7 ± 1.11 %, middle_Lmo3-/-_ = 21.8 ± 1.4 %, *p* = 0.0015; caudal_WT_ = 25.3 ± 1.1 %, caudal_Lmo3-/-_ = 17.7 ± 1.4 %; *p* = 0.0044; two-tailed *t* test, *n*=5 for WT and 4 for *Lmo3*-null for rostral and caudal and *n*=4 for both genotypes for middle; Fig. 4 *K*).

**Figure 4.**
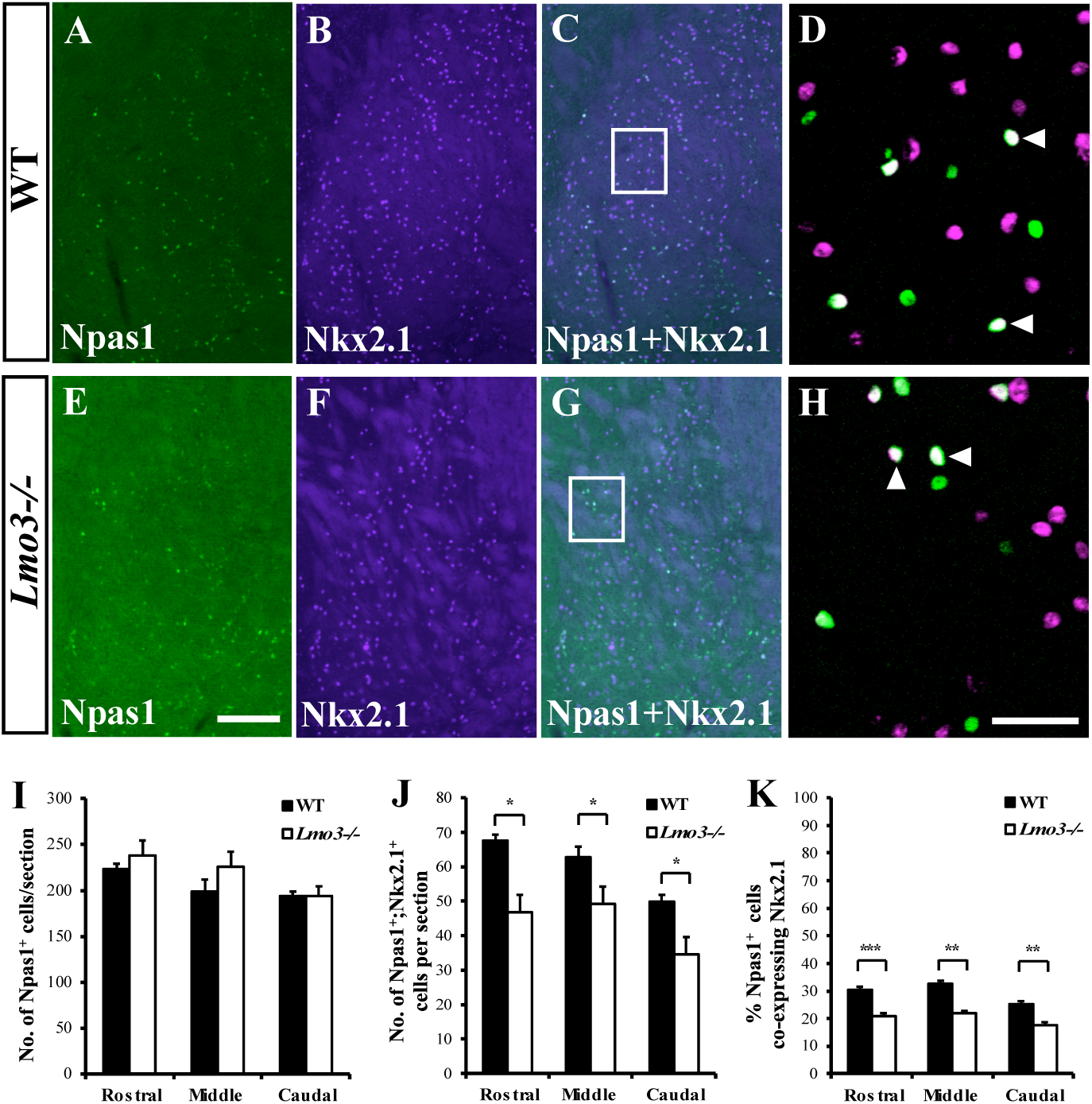
Npas1^+^;Nkx2.1^+^ neurons are reduced in the *Lmo3*-null mutant. Coronal views of the GPe of P30 wildtype (**A**-**C**) and *Lmo3*-null mutants (**E**-**G**) immunostained for Npas1 and Nkx2.1. **D** and **H** are high magnification images of the boxed regions in **C** and **G**, respectively. Filled arrowheads indicate Npas1^+^;Nkx2.1^+^ neurons. **I**-**K**, Quantification of Npas1^+^ (**I**), Npas1^+^;Nkx2.1^+^ (**J**) GPe neurons and % Npas1^+^ neurons co-expressing Nkx2.1^+^ (**K**) from three different rostrocaudal levels (see Fig. 2. For all graphs, *n*=4 or 5 for WT and *Lmo3-/-* for rostral and caudal levels and *n*=3 for WT and 4 for *Lmo3-/-* for the middle level). [Graphs indicate mean values; error bars show SEM. Two-tailed *t* test, \**p* < 0.05, \*\**p* < 0.01, \*\*\**p* < 0.001] Scale bars: (**E**) 200 μm; (**H**) 50 μm.

We could not directly assess a reduction in the neuronal subpopulation coexpressing Nkx2.1 and PV due to technical limitations (both antibodies were raised in mouse). Given that more than 80% of GPe PV^+^ neurons express Nkx2.1 and about two-thirds of Nkx2.1^+^ neurons express PV (Dodson et al., 2015; Abecassis et al., 2020), it is likely that the Nkx2.1^+^;PV^+^ subset is significantly reduced in the mutant. In addition, our findings indicate that a small proportion of Nkx2.1^+^ neurons that are reduced in the *Lmo3*-null belong to the Npas1^+^;Nkx2.1^+^ subset. While this may seem surprising given the overall unchanged number of Npas1^+^ neurons in the mutant, there appeared to be a trend toward an increase in FoxP2^+^ neurons (most of which also express Npas1^+^) in the mutant (Fig. 3 *I*,*J*,*K*), even though this was not statistically significant. The neuronal makeup of the GPe assessed using different molecular markers in the wildtype (Table 1) is consistent with values reported in prior studies from different laboratories (Nóbrega-Pereira et al., 2010; Dodson et al., 2015; Saunders et al., 2016; Abecassis et al., 2020). The percent of Npas1^+^ neurons expressing FoxP2, examined in a fewer number of sections, appeared slightly higher in the mutants (WT = 58.2 ± 6.4 %, *Lmo3-/-* = 64 ± 2.3 %, 5 sections from 2 brains for WT and 6 sections from 2 brains for *Lmo3*-null), as did the percent of Npas1^+^ neurons expressing PV (WT = 4 ± 0.4 %, *Lmo3-/-* = 12.1 ± 10 %, 4 sections from 2 brains for WT and 5 sections from 2 brains for *Lmo3*-null), though values for both WT and mutants fall within the range reported in earlier studies (Dodson et al., 2015; Hernández et al., 2015; Abrahao and Lovinger, 2018; Abecassis et al., 2020).

**Table 1.**

|  | PV<sup>+</sup> |  | Nkx2.1<sup>+</sup> |  | Npas1<sup>+</sup> |  |
| --- | --- | --- | --- | --- | --- | --- |
|  | % of total<br>GPe<br>neurons | *n* (mice) | % of total<br>GPe<br>neurons | *n* (mice) | % of total<br>GPe<br>neurons | *n* (mice) |
| WT | 43.1 ± 3.2 | 3 | 65 ± 2.6 | 2 | 34.8 ± 0.8 | 2 |
| *Lmo3*<sup>-/-</sup> | 30.4 ± 1.6 | 3 | 50.6 ± 0.7 | 2 | 46.2 ± 0.4 | 2 |

% values indicate mean ± SEM
NeuN was used as the marker for total GPe neuron counts per section.
For each *n* of all 3 molecular markers, counts were obtained from 2-3 sections per level (Fig. 2I).

### Loss of *Lmo3* results in a reduction of the pallidosubthalamic input

The reciprocally connected GPe and subthalamic nucleus (STN) network is integral to basal ganglia function and pallidal projections to the STN strongly influence and pattern the activity of STN neurons (Bevan et al., 2002; Baufreton et al., 2005, 2009; Atherton et al., 2013). In chronic mouse models of PD, maladaptive alterations in the strength of the GPe-STN input are thought to exacerbate the pathological firing observed in the disease (Fan et al., 2012; Chu et al., 2015). PV^+^ neurons constitute the major pallidal input to the STN (Hoover and Marshall, 1999; Mallet et al., 2012; Mastro et al., 2014; Abdi et al., 2015; Hernández et al., 2015; Abecassis et al., 2020; Pamukcu et al., 2020). Since these neurons are reduced in the *Lmo3*-null mutants, we wanted to determine if the pallidosubthalamic input is also reduced in the mutants. Pallidosubthalamic projections form the main GABAergic input to the STN (Canteras et al., 1990; Smith et al., 1998; Glajch et al., 2016). Thus, we performed immunostaining at P30 for vesicular GABA transporter (vGAT), a marker of GABAergic axon terminals (McIntire et al., 1997), and the presynaptic active zone protein Bassoon (Tom Dieck et al., 1998). Visual inspection of synaptic marker staining in the STN of mutant sections suggested a considerable reduction in size; thus we first quantified the area labeled by vGAT and Bassoon immunostaining in the STN, using borders marked by NeuN as an approximate boundary (Fig. 5 *A*-*D, A’*-*D’*). This revealed a significant reduction in area in the *Lmo3*-null mutants compared to wildtype (WT = 4.57 ± 0.24 x10^5^ µm^2^, *Lmo3-/-* = 3.42 ± 0.18 x10^5^ µm^2^, *p* = 0.0048, two-tailed *t* test, *n*=5 for both genotypes; Fig. 5 *E*). PV immunoreactivity in the STN was also markedly reduced in the *Lmo3*-null mutants (Fig. 5 *F*-*J, F’*-*J’*). This PV immunostaining primarily represents pallidosubthalamic axon terminals as *in situ hybridization* data indicates that PV^+^ STN neurons are found in a restricted, dorsolateral domain of the structure (Wallén-Mackenzie et al., 2020; Jeon et al., 2022), and cell body staining in the STN was not visible, unlike in the adjacent zona incerta (Fig. 5 *H,I’*). In addition, we employed a previously established approach (Fan et al., 2012; Chu et al., 2015) to analyze the density of inhibitory presynaptic structures in the STN, in order to assess if there were any changes in the *Lmo3*-null. From high magnification confocal images, we counted vGAT^+^ structures, as well as Bassoon^+^ structures that colocalized with vGAT^+^ structures, using an optical dissector method. However, this analysis did not reveal any changes in the density of vGAT^+^ or Bassoon^+^;vGAT^+^ structures in the *Lmo3*-null, as compared to WT (vGAT_WT_ = 41.02, 30.81 – 45.59 million/mm^3^, vGAT_Lmo3-/-_ = 37.51, 25.55 – 43.78 million/mm^3^, *p* = 0.193, Mann-Whitney *U* test, *n*=8 for both genotypes; and Bassoon;vGAT_WT_ = 44.2, 30.42 – 69.16 million/mm^3^, Bassoon;vGAT_Lmo3-/-_ = 37.05, 27.73 – 55.51 million/mm^3^, *p* = 0.5221, Mann-Whitney *U* test, *n*=7 for both genotypes; Fig. 5 *K*, *L*, *K’*, *L’*, *M*). The median and range of values for vGAT^+^ structures in the wildtype is consistent with an earlier study (Chu et al., 2015), although our values for Bassoon^+^;vGAT^+^ structures is slightly lower compared to the same, which could be attributed to the differences in age and sex of animals used (juvenile females and males in our study as compared to adult males in the earlier one).

**Figure 5.**
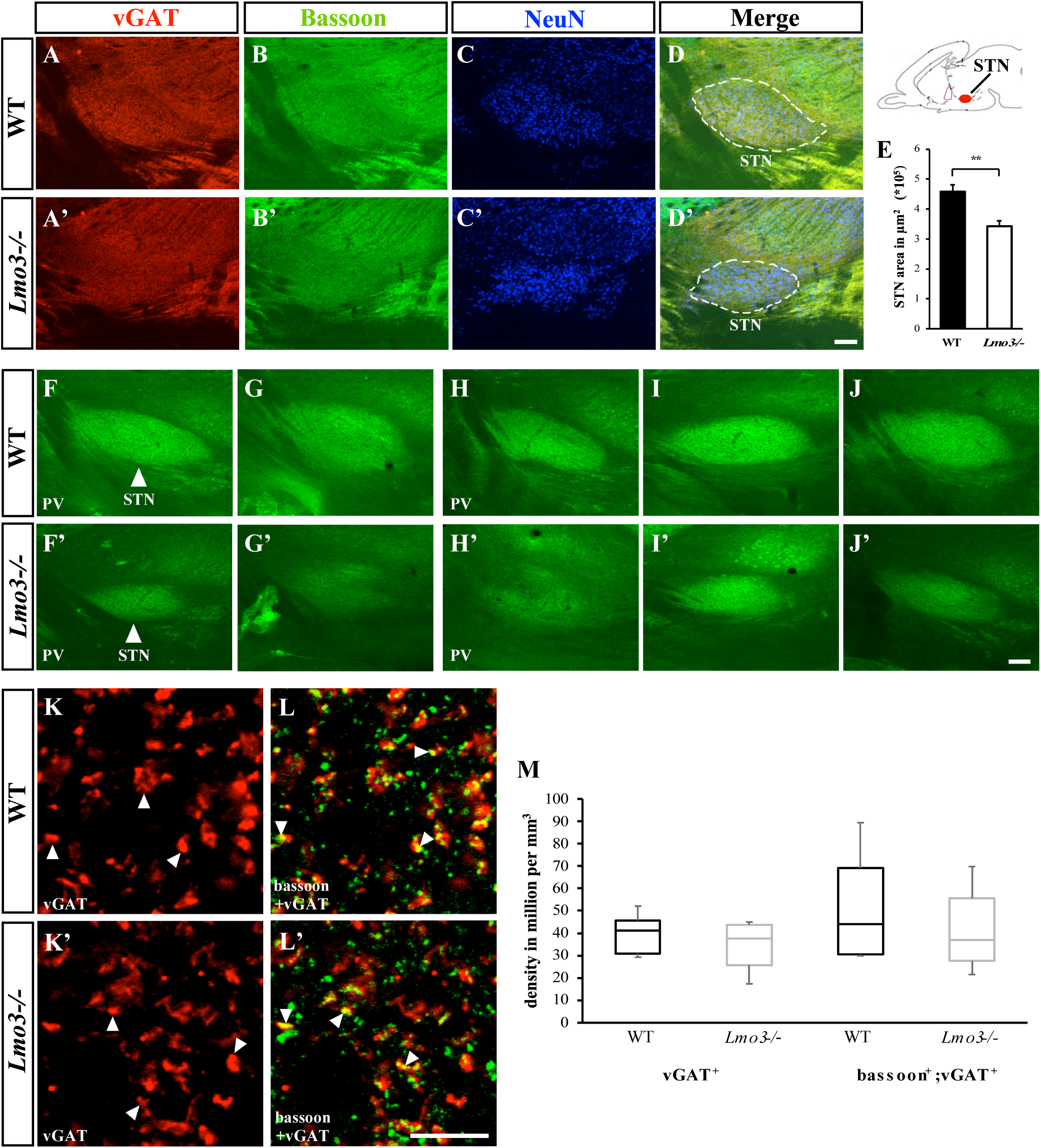
Loss of Lmo3 results in a reduction of the pallidosubthalamic input. Immunostaining for the presynaptic markers vGAT (**A**,**A’**) bassoon (**B**,**B’**) and NeuN (**C**,**C’**) in sagittal sections of subthalamic nucleus (STN) at similar medio lateral levels of wildtype and *Lmo3*-null mutants. E, Quantification of size of inhibitory input using NeuN to demarcate the STN (dashed outlines in **D**,**D’**), reveals a significant reduction in the *Lmo3*-null mutants (**E**), *n*=5 for WT and *Lmo3-/-* [Graph indicates mean values; error bars show SEM. Two-tailed *t* test, *p* = 0.0048] **F**-**J**, **F’**-**J’** show STN stained with PV antibody. **F**,**G** and **F’**,**G’** are sections from one mouse each for WT and *Lmo3-/-* and **H**-**J**, **H’**-**J’**, are from a second mouse. Arrowheads indicate the STN, which is smaller in size in the *Lmo3*-null mutants. **K**,**K’**, are representative confocal micrographs of vGAT immunostaining in the STN of WT and *Lmo3-/-* mice, respectively; **L**,**L’** demonstrate bassoon double immunostaining. Filled arrowheads indicate examples of vGAT^+^ and bassoon^+^;vGAT^+^ structures counted. **M**, Box-and-whisker plots showing density of vGAT^+^ (left) and bassoon^+^;vGAT^+^ structures (right) in the STN of WT and *Lmo3-/-* mice. Boxes denote first, second (median) and third quartiles, while whiskers represent the 10^th^ and 90^th^ percentiles. *n*=8 for WT and Lmo3-/- for quantification of vGAT^+^ and *n*=7 for quantification of bassoon^+^;vGAT^+^ structures. [Mann-Whitney *U* test, *p* = 0.193 for vGAT^+^ and *p* = 0.5221 for bassoon^+^;vGAT^+^] Scale bars: (**D’**,**J’**) 100 μm; (**L’**) 5 μm.

### Loss of *Lmo3* does not affect the development of cholinergic GPe neurons and striatal projection neurons

*Lmo3* is expressed broadly in regions of the developing subpallium (Fig. 1) that are the source of several basal ganglia neuronal subtypes, in addition to GABAergic GPe projection neurons. Cholinergic neurons form a relatively small proportion of the GPe, but project to the frontal cortex, thus forming a direct link between the basal ganglia and the cortex (Saunders et al., 2015; Abecassis et al., 2020). These cholinergic neurons arise primarily from the MGE (Flandin et al., 2010; Nóbrega-Pereira et al., 2010). We examined the *Lmo3*-null mutants for changes in the development of cholinergic neurons by performing immunostaining for choline acetyl transferase (ChAT) and found no significant differences between controls and *Lmo3*-null mutants in ChAT^+^ neurons along the border of the GPe and the nucleus basalis (Fig. 1-1 *A*, *A’*, *G*). Other cholinergic neuron subtypes: striatal ChAT^+^ interneurons and ChAT^+^ projection neurons in the septum and diagonal band that also arise from the MGE (Olsson et al., 1998; Marín et al., 2000; Zhao et al., 2003), were not affected either, in the *Lmo3*-nulls: see (Fig. 1-1 *B*,*C*,*D*,*E, B’*,*C’*,*D’*,*E’, H*) for striatal ChAT^+^ interneurons, and (Fig. 1-1 *F, F’*) for ChAT^+^ projection neurons in the septum and diagonal band. The spiny projection neurons (SPNs) which comprise the bulk of the striatum and form the principal inhibitory input to the GPe (Kita, 2007; Gerfen and Surmeier, 2011; Hegeman et al., 2016), are born in the LGE (Olsson et al., 1998; Wichterle et al., 2001). Immunostaining for dopamine- and cAMP-regulated neuronal phosphoprotein 32 (DARPP-32), a marker for all striatal projection neurons (Anderson and Reiner, 1991), revealed no gross changes in the development of SPNs of the striatum as judged by the normal innervation of targets by projection neurons of both the direct and indirect pathway (Fig. 1-1 *I*-*K, I’*-*K’*).

### Reduction of MGE derived neurons in the *Lmo3* null GPe is evident at birth

We next sought to determine whether we could observe the neuronal subtype specific changes seen in the GPe of mature *Lmo3*-null mice, at earlier time points. Since *Lmo3* is strongly expressed in the SVZ of the MGE as well as the developing GPe during embryonic time points when GPe neurons are born (Fig. 1) and (Marchand and Lajoie, 1986; Nóbrega-Pereira et al., 2010), we hypothesized that Lmo3 is required during embryonic time points for correct cell fate specification in the GPe. We thus analyzed the control and *Lmo3*-null GPe at P0. Since GPe neurons do not express detectable levels of PV till later postnatal stages (Mitrofanis, 1992), we performed *in situ* RNA hybridization for Er81, a transcription factor that is expressed in most PV^+^ neurons in the adult GPe (Nóbrega-Pereira et al., 2010) and found that *Er81* expression is strongly reduced in the GPe of P0 *Lmo3*-null mice as compared to controls (Fig. 6 *I*-*N*). In addition, we also performed immunohistochemistry for Nkx2.1 and FoxP2 (Fig. 6 *A*-*F*). In the mutant, Nkx2.1^+^ neuron number was reduced (rostral_WT_ = 738 ± 2, rostral_Lmo3-/-_ = 500.9 ± 13.4, *p* = 0.0032; caudal_WT_ = 431.1 ± 48.7, caudal_Lmo3-/-_ = 203.4 ± 22.3; *p* = 0.03; two-tailed *t* test, *n*=3 for both genotypes; Fig. 6 *H*) whereas the numbers of FoxP2^+^ neurons were unaltered compared to control (rostral_WT_ = 231.3 ± 28.3, rostral_Lmo3-/-_ = 253.6 ± 46.4, *p* = 0.709; caudal_WT_ = 221.6 ± 12.4, caudal_Lmo3-/-_ = 267.7 ± 33.4; *p* = 0.324; two-tailed *t* test, *n*=3 for both genotypes; Fig. 6 *H*) The number of Nkx2.1^+^ neurons in both the WT and *Lmo3*-null is higher at P0 than at P30 (compare Fig. 3 *I*,*J*,*K* to Fig. 6 *H*), which is likely due to the fact that Npas1^+^ neurons progressively downregulate Nkx2.1 expression postnatally (Flandin et al., 2010; Abdi et al., 2015; Dodson et al., 2015; Abecassis et al., 2020).

**Figure 6.**
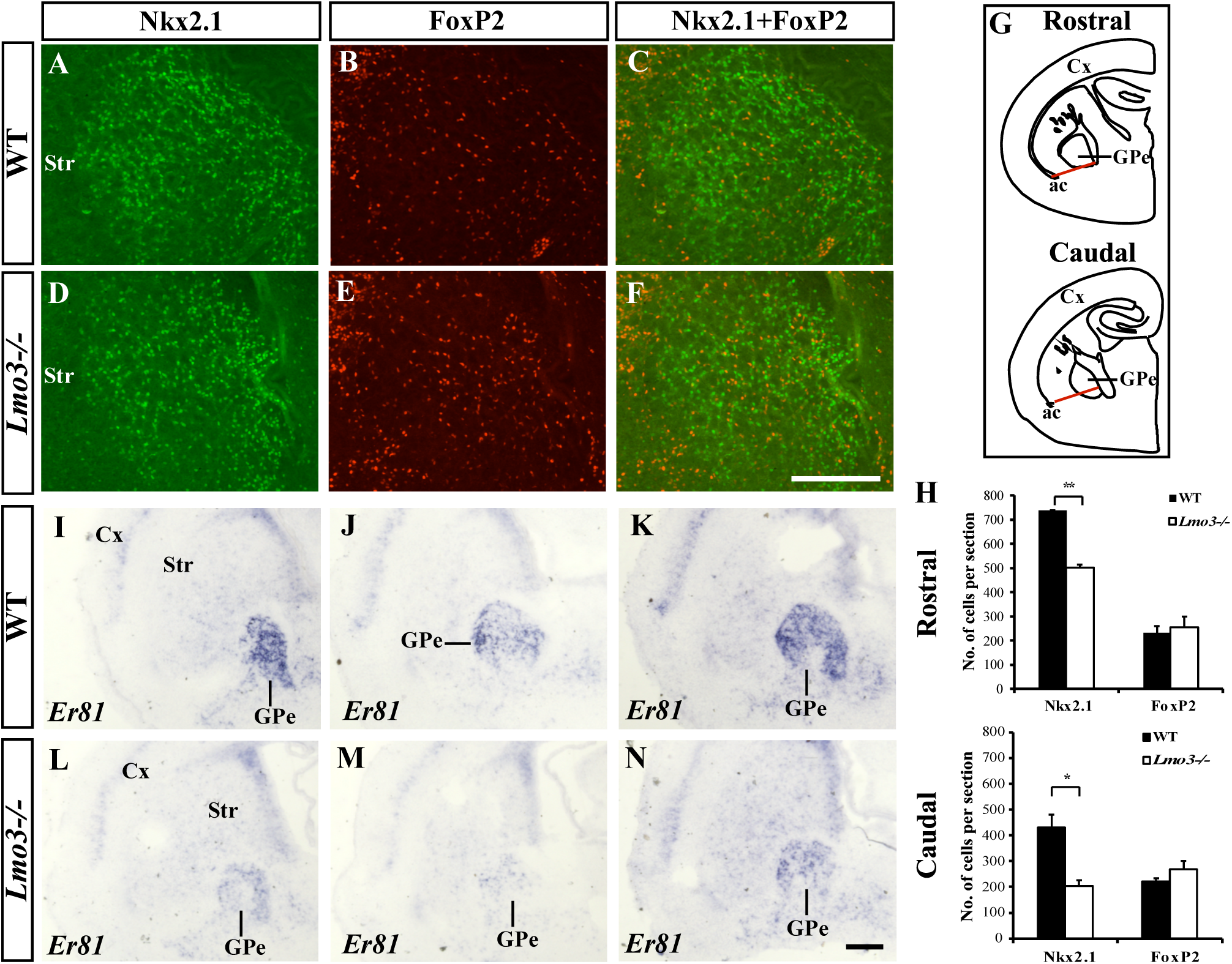
Reduction of MGE derived GPe neurons in the *Lmo3-*null mutant at P0. **A**-**F**, Immunohistochemistry for Nkx2.1 and FoxP2 in the GPe of wildtype (**A**-**C**) and *Lmo3*-null (**D**-**F**) mutants. **G**, Coronal sections illustrating 2 different rostrocaudal levels of the GPe from which neurons were counted. Ventral borders of the GPe (red lines) were defined according to the medial edge of the anterior commissure (ac). **H**, Quantification shows a significant reduction of Nkx2.1^+^ neurons in the *Lmo3-/-* GPe at both rostrocaudal levels. No significant differences in FoxP^+^ neuron numbers were observed between *Lmo3-/-* and WT (*n* = 3 each for WT and *Lmo3-/-* for both graphs). [Graphs indicate mean values; error bars show SEM. Two-tailed *t* test, At the rostral level, *p*= 0.003 for Nkx2.1 and 0.709 for FoxP2; for caudal level *p* = 0.035 for Nkx2.1 and 0.324 for FoxP2. \**p* < 0.05, \*\**p* < 0.01] **I**-**N**, *In situ* RNA hybridization staining at three different rostrocaudal levels shows strong *Er81* expression labeling the P0 GPe in WT (**I**-**K**), and a marked reduction in the *Lmo3-/-* GPe (**L**-**N**). Cx, cortex; GPe, external globus pallidus; Str, striatum. Scale bars: 200 μm.

### Analysis of *Lmo3* null mice at embryonic time points

The fact that Nkx2.1^+^ neurons and *Er81* expression in the *Lmo3* null mutants are already reduced at P0 supports the idea that *Lmo3* is required during prenatal stages for development of GPe neurons. To assess whether reduced proliferation may contribute to this phenotype, we assessed phospho-histone 3 (pH3) expressing (M-phase) cells in the MGE of E11.5, E12.5 and E13.5 controls and Lmo3 mutants. We calculated the density of pH3^+^ cells in both the ventricular zone and subventricular zone of the ventral MGE which is the source of PV^+^ and Nkx2.1^+^ GPe neurons (Flandin et al., 2010; Nóbrega-Pereira et al., 2010), but found no significant difference between controls and Lmo3 mutants in either the caudal or rostral MGE (See Fig. 6-1).

GPe development has been shown to depend on transcription factors that control ventral telencephalon patterning and differentiation, like Dlx2 and Mash1 (Long et al., 2009a, 2009b; Nóbrega-Pereira et al., 2010), and Nkx2.1, which is a primary determinant of MGE identity is also required at later stages of development of MGE derived neurons like cortical and striatal interneurons (Sussel et al., 1999; Du et al., 2008; Butt et al., 2008). We thus analyzed the expression of these genes in the *Lmo3* mutants at E13.5. However, as shown in Fig. 7-1, loss of *Lmo3* does not alter the expression of these transcription factors. Interestingly, there appears to be a mild reduction of *Lmo1* expression in the *Lmo3*-null (Fig. 7-1 *J*, *K*, *L*, *J’*, *K’*, *L’*). The LIM-HD gene *Lhx6*, a transcriptional target of Nkx2.1 (Du et al., 2008), is required for the differentiation, but not subtype specification, of MGE derived cortical interneurons (Liodis et al., 2007; Du et al., 2008), while *Lhx6* and *Lhx8* double mutants have defects in GPe development (Flandin et al., 2011). Expression of these genes was also unchanged in the *Lmo3*-null mutants (Fig. 7-2), which is in line with the fact that they are likely upstream of Lmo3 in development (Flandin et al., 2011). In addition, the *Lmo3*-null mutant displayed no changes in the expression patterns of *Lmo4* and *Pax6* (Fig. 7-2), which are markers for the LGE (Flandin et al., 2010; Long et al., 2009b; Stoykova et al., 2000).

Both PV expressing GPe projection neurons and PV expressing cortical interneurons arise from the MGE, although PV^+^ cortical interneurons are generated over a more protracted time frame as compared to PV^+^ GPe neurons (Miyoshi et al., 2007; Nóbrega-Pereira et al., 2010; Inan et al., 2012). In addition, PV^+^ GPe neurons arise from a more caudoventral region of the MGE as compared to PV^+^ cortical interneurons (Flandin et al., 2010; Inan et al., 2012; Nóbrega-Pereira et al., 2010). Since *Lmo3*-null mutants also have reduced numbers of PV^+^ cortical interneurons (Au et al., 2013), we next analyzed genes which have gradients of expression along the rostrocaudal and dorsoventral axes of the MGE, and/or are involved in specification of MGE derived cortical interneuron subtypes, such as *Er81*, *Couptf1* and the Sonic Hedgehog (Shh) signaling effectors Gli1 and Ptc1 (Flames et al., 2007; Wonders et al., 2008; Nóbrega-Pereira et al., 2010; Xu et al., 2010; Flandin et al., 2011; Lodato et al., 2011; Hu et al., 2017). Expression patterns of *Er81*, *Couptf1*, *Gli1* and *Ptc1* appeared to be unaltered in the ventral telencephalon of the *Lmo3*-null mutants (Fig. 7). Expression patterns of several of the above genes (*Dlx2, Couptf1, Er81, Pax6*) were also analyzed at E12.5, with similar results as above.

**Figure 7.**
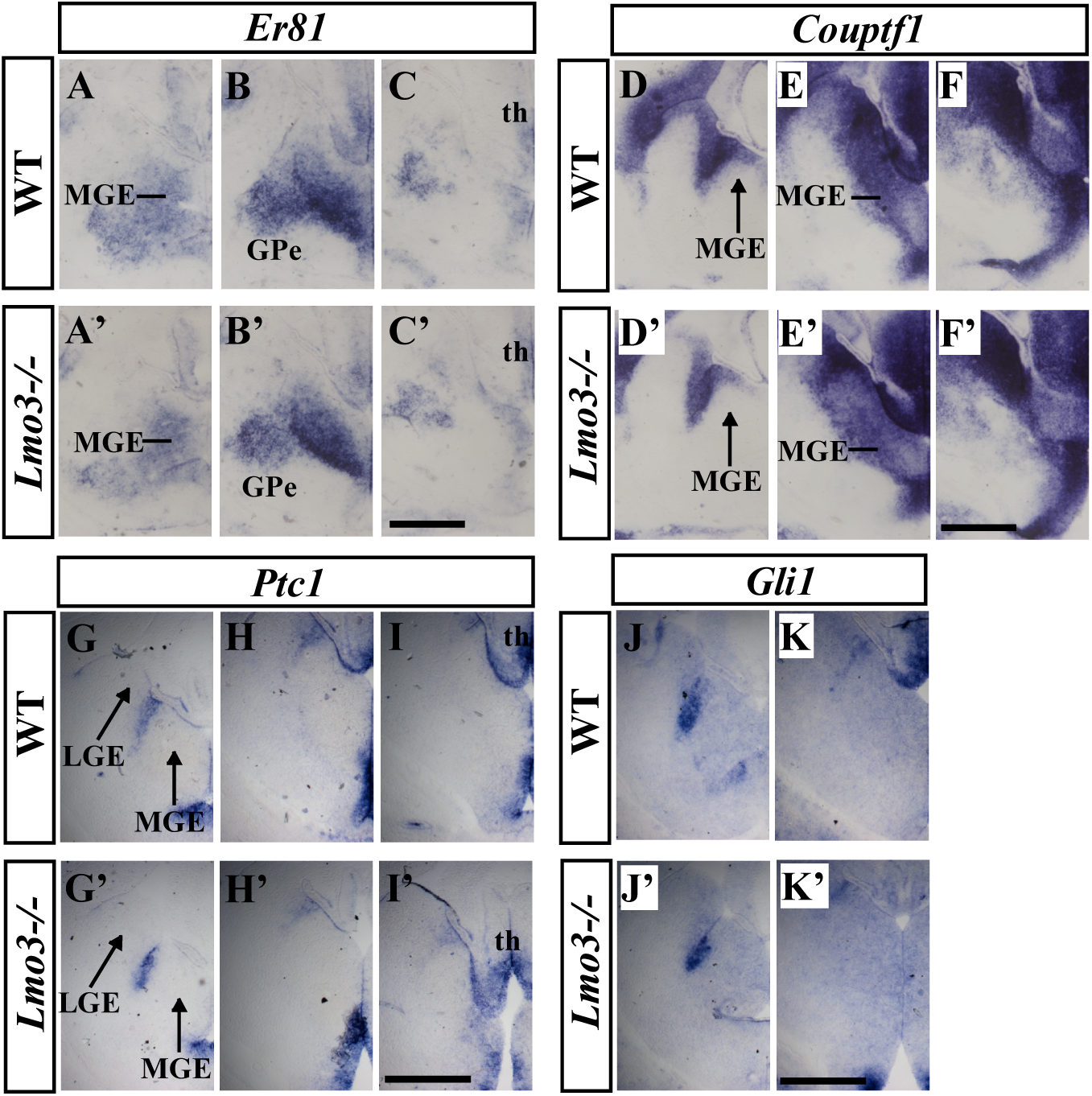
Expression of selected MGE markers is unchanged in the *Lmo3-*null mutant at E13.5. Images of three serial coronal hemisections (rostral-most on the left) show *in situ* RNA hybridization staining for *Er81* (**A**-**C**, **A’**-**C’**), *Couptf1* (**D**-**F**, **D’**-**F’**), *Ptc1* (**G**-**I**, **G’**-**I’**) and *Gli1* (**J**,**K**,**J’**,**L’**). Cx, cortex; GPe, external globus pallidus; LGE, lateral ganglionic eminence; MGE, medial ganglionic eminence; th, thalamus. Scale bars: 500 μm.

## Discussion

Studies over the past decade have delineated different subtypes of GABAergic GPe projection neurons based on molecular characteristics, connectivity patterns and functional criteria, which has improved our understanding of the role of the GPe in regulating motor function and dysfunction (Mallet et al., 2012; Abdi et al., 2015; Dodson et al., 2015; Hernández et al., 2015; Mallet et al., 2016; Abrahao and Lovinger, 2018; Abecassis et al., 2020; Pamukcu et al., 2020; Aristieta et al., 2021; Ketzef and Silberberg, 2021; Cui et al., 2021a). Despite this, little remains known about the molecular mechanisms that underlie the development of these GPe neuronal subtypes. Our results demonstrate that Lmo3 is required for the development of MGE derived PV^+^ and Nkx2.1^+^ GPe neurons, but not FoxP2^+^ neurons, Npas1^+^ neurons overall, or cholinergic neurons. Development of striatal projection neurons appears normal in *Lmo3*-null mutants. As a consequence of the reduction in PV^+^ neurons, *Lmo3*-null mice have a reduced pallidal input to the subthalamic nucleus.

There can be differences within a GPe neuronal subpopulation expressing the same molecular marker. For example, the use of transgenic reporter lines driven by subpallial enhancer elements has shown that there are at least two pools of PV^+^ neuron progenitors in the MGE: one that generates predominantly rostral PV^+^ neurons, and the other, rostral and caudal neurons but with a bias to the latter (Silberberg et al., 2016). In agreement with this, single-cell transcriptomics analysis confirms the existence of two PV^+^ neuron clusters within the GPe (Saunders et al., 2018), while a recent study suggested subtle differences in behavior controlled by the two PV^+^ neuronal subtypes belonging to these two clusters (Cui et al., 2021b). In light of these observations, it is interesting that the loss of Lmo3 results in a more severe reduction of caudal GPe PV^+^ neurons than at rostral and middle levels (Fig. 2 *J*,*K*,*L*). The reduction of Npas1^+^;Nkx2.1^+^ neurons upon loss of Lmo3 is consistent with the rest of our data, in that the development of MGE but not LGE derived GABAergic neurons is affected in the mutants, in spite of Lmo3 being expressed in both the MGE and LGE. Although Npas1^+^;Nkx2.1^+^ neurons form a small proportion (∼10%) of GPe neurons, they constitute the non-cholinergic GPe projection to the cortex; densely innervating deep layers of the primary motor and somatosensory cortices (Abecassis et al., 2020). Lhx6 is a marker for another subpopulation of GPe neurons derived from the MGE (Dodson et al., 2015). Different subpopulations of Lhx6^+^ GPe neurons coexpress either PV or Npas1; although there is another, unique subpopulation of Lhx6^+^ neurons that expresses neither PV nor Npas1 (Mastro et al., 2014; Dodson et al., 2015; Hernández et al., 2015; Abrahao and Lovinger, 2018; Abecassis et al., 2020). Lhx6^+^ neurons have electrophysiological properties typically in between the extremes of PV^+^ and FoxP2^+^ neurons, with projections to other basal ganglia structures more similar to PV^+^ neurons than FoxP2^+^ neurons (Mastro et al., 2014; Hernández et al., 2015; Abrahao and Lovinger, 2018; Abecassis et al., 2020). Lhx6 expression does not appear markedly changed at a later embryonic age (E15.5) or at P0 in the GPe of *Lmo3*-null mutants: supplementary data in (Au et al., 2013) and Fig. 7-3. It would be interesting to address whether Lhx6^+^;PV^+^, Lhx6^+^;Npas1^+^ and Lhx6^+^;PV^-^; Npas1^-^ GPe neurons are spared upon loss of Lmo3 in future experiments, employing the transgenic mouse line used in the studies mentioned above, for consistency.

While we have shown that loss of Lmo3 results in a reduction of MGE derived GPe neurons by birth, analysis of the *Lmo3*-null mutants at embryonic ages did not reveal any obvious underlying mechanism for the cell subtype specific requirement for Lmo3 in their development. Expression of transcription factors implicated in earlier stages of ventral telencephalon patterning, like *Dlx2*, *Mash1*, *Nkx2.1*, were unaffected in the *Lmo3*-null. The expression of several genes that are expressed in gradients along the rostrocaudal and dorsoventral axes of the MGE (*Couptf1, Gli1, Ptc1*, *Er81)*, and genes that have been shown to influence the development of PV^+^ expressing neurons (*Couptf1, Gli1, Ptc1)* (Flames et al., 2007; Xu et al., 2010; Lodato et al., 2011) was also unaffected in the *Lmo3*-null mutant.

Our data show a significant reduction of area innervated by pallidal axon terminals in the STN of *Lmo3*-null mutants. Similar to this, an earlier study that had investigated the role of the transcription factor Isl1 (which belongs to the same protein family as Lmo3) in the development of striatonigral neurons, showed that striatonigral neurons are reduced in *Isl1* knockout mutants, leading to a considerable reduction in their area of innervation of the EP and SNr (Ehrman et al., 2013). We performed the quantitation of synaptic marker density from roughly the center of the STN in all sections analyzed in both genotypes, which did not demonstrate any significant change in the *Lmo3*-null, as compared to WT. Nonetheless, an aspect of future work could be functional experiments to determine if GPe-STN synaptic transmission is altered in *Lmo3*-null mutants, and at adult ages in addition to juvenile time points.

Collectively, recent studies indicate that a major function of PV^+^ neurons and many Nkx2.1^+^ neurons, is what was traditionally ascribed to the GPe as a whole. As part of the indirect pathway of the basal ganglia, they receive strong inhibitory input from indirect pathway striatal projection neurons (iSPNs) (Aristieta et al., 2021; Ketzef and Silberberg, 2021; Cui et al., 2021a); a decrease in their firing would disinhibit the STN as well as basal ganglia output nuclei (SNr) in which the axons of many PV^+^ and Nkx2.1^+^ neurons also collateralize, thus suppressing movement (Mallet et al., 2012; Sano et al., 2013; Mastro et al., 2014); and optogenetic activation of PV^+^ neurons and their terminals in the STN has been shown to promote movement (Pamukcu et al., 2020). The firing of FoxP2^+^ and Npas1^+^ neurons would depend on how they integrate input from various sources: motor cortex, dSPNs, iSPNs, STN as well as from PV^+^ neurons (Mallet et al., 2012; Karube et al., 2019; Aristieta et al., 2021; Ketzef and Silberberg, 2021; Cui et al., 2021a). As regards their role in locomotor behavior, activation of FoxP2^+^ neurons has been linked to cancellation of ongoing or imminent action (Mallet et al., 2016; Aristieta et al., 2021) while activation of Npas1^+^ neurons as well as FoxP2^+^ neurons decreased movement speed and increased the length of time mice spent motionless (Glajch et al., 2016; Pamukcu et al., 2020; Cui et al., 2021b). These behaviors are likely mediated through the dense axonal projections of these neurons to the striatum, though it remains to be determined which of the striatal neuronal subtypes they innervate are involved in specific aspects of their motor control, and if additional neuronal circuits are involved. Given the above, we could speculate that the reduced PV^+^ and Nkx2.1^+^ neurons and pallidosubthalamic input in *Lmo3*-null mice, along with potentially reduced PV^+^ and Nkx2.1^+^ inhibitory input to FoxP2^+^ neurons whose development remains unaffected, would result in reduced motor activity. Data from two studies support this hypothesis: overall locomotor activity was shown to be reduced in *Lmo3*-null mice compared with wildtype controls, and this effect could not be attributed to the function of Lmo3 in the brain regions and behaviors under investigation-i.e., amygdala and anxiety-like behavior (Savarese and Lasek, 2018; Reisinger et al., 2020).

A recent study that used a novel deep brain stimulation (DBS) protocol to drive neuronal subtype-specific neuromodulation in the GPe by harnessing differences in the properties of PV^+^ and Lhx6^+^ GPe neurons, achieved longer lasting therapeutic effects than that obtained using a conventional DBS protocol, in an acute 6-OHDA lesioned mouse model of PD (Spix et al., 2021). Such progress underlines the importance of understanding the organization of basal ganglia structures, including molecular factors that control their development.

## Materials and Methods

### Animals

The Lmo3-LacZ knock-in construct was generated in the laboratory of Dr Terence Rabbitts, University of Oxford. Briefly, a *LacZ* gene expressed from an EMC-IRES promoter was inserted into exon 2 of the *Lmo3* gene to interrupt the *Lmo3* coding region and generate a *Lmo3* loss-of-function allele (Tse et al., 2004). Targeted ES cells were obtained from Rabbitts. Clones of these targeted ES cells were injected into C57BL/6J blastocysts to generate mouse chimeras in our laboratory. Heterozygous *Lmo3^LacZ/+^* mice were generated in 129S6 and C57BL/6J mixed background as described before (Gan et al., 1996, 1999). The PCR primers used to identify the *Lmo3-LacZ* knock-in allele were 5’-CGAGTTCTTCAGTCAGAGTAC-3’ and 5’-TCCTGCCCAGAAAGTATCCAT-3’. PCR primers used to identify the Lmo3 wildtype allele were 5’-TGCTGTCAGTTCAGCCAGAC-3’ and 5’-CGAGTTCTTCAGTCAGAGTAC-3’. The *Lmo3^LacZ/LacZ^* homozygous mice are referred to as *Lmo3*-null or *Lmo3-/-* in the rest of the paper. *Lmo3^LacZ/+^* mice are referred to as *Lmo3+/-*, or heterozygous mice. *Lmo3*-null mice occasionally appeared a bit smaller in size and lighter in weight than their heterozygous and wildtype littermates but were capable of normal breeding and did not display increased mortality. We did not observe any obvious difference between the controls and mutants regarding the size of the forebrain. For the staging of embryos and adult mice, noon of the day vaginal plugs were observed was considered E0.5. For the P0 time point, pups were harvested within 24 hours of birth. All animal procedures in this study were in accordance with NIH guidelines and were approved by the University Committee of Animal Resources (UCAR) at the University of Rochester.

### Tissue preparation

For embryonic ages and P0, decapitated heads were drop fixed in 4% paraformaldehyde (PFA) for 8-48 hours at 4°C. P30 and older mice were transcardially perfused with 4% PFA followed by 30 −60 minutes postfixation at 4°C. Following postfixation, tissue was equilibrated in 30% sucrose at 4°C until it sunk to the bottom. Tissue was frozen in OCT compound (Tissue-Tek) (embryonic heads and P0 brains), or TFM medium (Electron Microscopy Sciences) (P30 brains) and stored at −80°C. Embryonic and P0 tissue was cryosectioned between 20-25 μm and collected on glass slides (Fisher) for processing while P30 and older brains cryosectioned at 40 μm and processed free-floating.

### Immunohistochemistry

Sections were blocked with 10% (v/v) normal goat serum (Invitrogen) and 0.1% (v/v) Triton X-100 in PBS for 30–60 min at 4°C and were subsequently incubated in primary antibodies in 1% normal goat serum and 0.1% Triton X-100 in PBS for 24 - 48 h at 4°C. After washes in PBS, the sections were incubated with Alexa Fluor-conjugated secondary antibodies in 0.1% Triton X-100 in PBS at room temperature for 2 h. For the phosphoHistone 3 (pH3) (embryonic time points E11.5, E12.5, E13.5) and Nkx2.1+FoxP2 (at P0) immunohistochemistry, antigen retrieval by heat pre-treatment (in sodium citrate buffer, pH6) was carried out prior to the blocking step. The primary antibodies and dilutions used in this study were: mouse anti-parvalbumin (1:250; Sigma P3088), mouse anti-TTF1 (Nkx2.1) (1:100; Novocastra; Leica Biosystems NCL-L-TTF-1), mouse anti-NeuN (1:500; Millipore MAB377), rabbit anti-NeuN (1:500; Millipore ABN78), goat anti-FoxP2 (1:500; Santa Cruz Biotechnology sc-21069), mouse anti-vGAT (1:1000; Synaptic Systems 131011), guinea pig anti-Bassoon (1:500; Synaptic Systems 141004), goat anti-ChAT (1:200; Millipore AB144P), rabbit anti-DARPP-32 (1:1000; Millipore AB10518) and rabbit anti-phosphoHistone 3 (1:250; Santa Cruz Biotechnology). Alexa Fluor conjugated secondary antibodies (Molecular probes) were used at a concentration of 1:1000 for all experiments except for synaptic marker (vGAT and Bassoon) immunostaining, for which they were used at a concentration of 1:500.

### *in situ* hybridization

*in situ* hybridization was performed on 20-35 μm thick sections using digoxigenin-labeled riboprobes, as described previously (Bulchand et al., 2003). The following probes were used: Lmo1, Lmo3, Lmo4, Mash1, Pax6, Isl1. Couptf1, Er81, Nkx2.1, Dlx2, Lhx6, Lhx8, Gli1, Ptc1 (gifted by Gord Fishell).

### Image acquisition and data analysis

Images of *in situ* hybridization experiments were captured using brightfield. For the analysis of pH3 and ChAT immunostaining and to obtain other low magnification immunofluorescence images, Axio Imager M1 (Carl Zeiss) was used. Z stack images were obtained using a confocal microscope LSM 510 (Carl Zeiss). Z stack images were obtained with optical slices 4 μm apart using the 40x water immersion objective for analysis of P0 GPe immunostaining. Cells were counted from the entire GPe, using the ventral borders defined according to the medial edge of the anterior commissure (Fig. 2*I*) 3 GPe sections were analyzed for each rostrocaudal level per brain. For ∼P30 (between postnatal days P30-P32) GPe immunostaining (Figs. 2, 3 and 4), Z stacks were obtained with optical slices 4 μm apart using the 20x objective. Cells were counted from the entire GPe. Ventral borders were defined according to the medial edge of the anterior commissure in rostral and middle sections and drawn at the same approximate dorsoventral level as the reticular nucleus of the thalamus for caudal sections (Fig. 2*I*). Between 3-4 sections were analyzed for each GPe rostrocaudal level per brain. For immunostaining with multiple markers, cells were counted in individual channels before quantifying dual labeling in merged images.

For STN area quantification at P30 (Fig. 5*E*), 4 different STN sections between ∼1.4 and 1.8 mm lateral of Bregma were analyzed per brain. For confocal analysis of bassoon and vGAT immunostaining, Z stack images were obtained at a thickness of 0.4 μm using the 100x oil immersion objective. Density of vGAT^+^ structures and bassoon^+^/vGAT^+^ structures was assessed using the optical dissector method (West, 1999., Fan et al., 2012). Sample sites were chosen using a grid (frame size, 10 μm x 10 μm) that was superimposed randomly on each image stack. Stereological counting was done through 4.4 μm thickness of the Z-stack (commencing at an optical section ∼ 4.5 μm below the slice surface). Immunoreactive structures were counted if they appeared within the sample frame and in the reference but not the adjacent look up optical section. vGAT^+^ structures > 0.5 μm^2^ in their maximal cross-sectional area were first selected and then bassoon^+^ structures that co-localized with them were subsequently counted.

Analyses were performed blind to genotype, using Image J (NIH). Statistical analyses were performed with Prism6 (GraphPad) and Microsoft Office Excel. All sample sizes (*n* values) reported represent number of mice.

## Acknowledgements

This work was supported by The National Institutes of Health EY026614 to L.G., the Research to Prevent Blindness challenge grant to the Department of Ophthalmology at the University of Rochester, NIMH R01 MH081880 to J.L.R.R. and NIH R01 NS069777 to C.S.C.

## Extended Data

**Figure 1-1.**
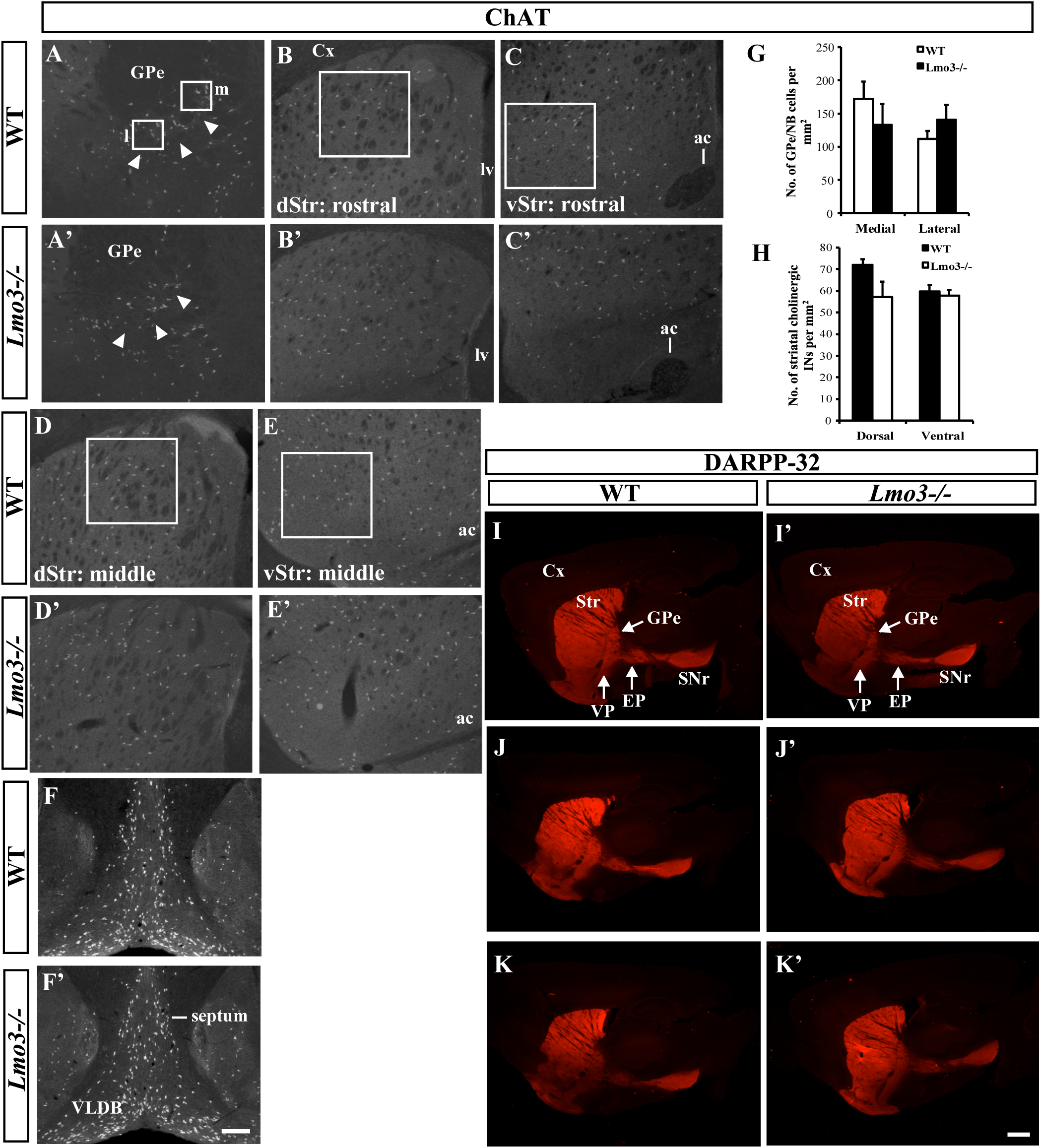
Lmo3 is not required for the development of cholinergic neurons and striatal projection neurons. **A**,**A’**, ChAT immunostaining for cholinergic neurons in the GPe and immediately adjacent nucleus basalis; the latter indicated by arrowheads. and the septum and VLDB. **B**,**B’**,**C**,**C’**, cholinergic interneurons at a rostral level of the striatum. **D**,**D’**,**E**,**E’**, cholinergic interneurons at a mid-level of the striatum. **F**,**F’**, cholinergic neurons in the septum and VLDB. Representative examples of regions from which cell counts were obtained are indicated by the white boxes in **A**,**B**,**C**,**D**,**E**. **G**, ChAT^+^ cell counts obtained from medial (m) and lateral (l) regions of the GPe-nucleus basalis boundary as indicated in (**A**), show no significant changes in the *Lmo3-*null: medial_WT_ = 172.5 ± 25.5, lateral_WT_ = 112 ± 11.5; medial_Lmo3-/-_ = 132.7 ± 31.4, lateral_Lmo3-/-_ = 140.9 ± 22.4, two-tailed *t* test, *p* = 0.381 for medial and *p* =0.315 for lateral. **H**, ChAT^+^ cell counts obtained from dorsal and ventral striatal regions of both rostral and middle levels of the striatum show no significant changes in the *Lmo3-*null: dorsal_WT_ = 72 ± 2.4, ventral_WT_ = 59.6 ± 3.1; dorsal_Lmo3-/-_ = 57.2 ± 7.1, ventral_Lmo3-/-_ = 57.7 ± 2.7, two-tailed *t* test, *p* = 0.161 for dorsal and *p* =0.667 for ventral. [Graphs indicate mean values; error bars show SEM. n=3 each for controls and mutants in both graphs]. Immunostaining for DARPP-32 at various medio-lateral levels in the WT (**I**-**K**) and *Lmo3-null* mutant (**I’**-**K’**) at P30 reveals no obvious changes in direct pathway projections (see EP and SNr) and indirect pathway projections (see GPe and VP). ac, anterior commissure, Cx, cortex; EP, entopeduncular nucleus; dStr, dorsal striatum; GPe, external globus pallidus; lv, lateral ventricle; NB, nucleus basalis; Str, striatum; SNr, substantia nigra pars reticulata; VP, ventral pallidum; VLDB, ventral limb of the nucleus of the diagonal band; vStr, ventral striatum. Scale bars: (**F’**) 200 μm; (**K’**) 500 μm.

**Figure 6-1.**
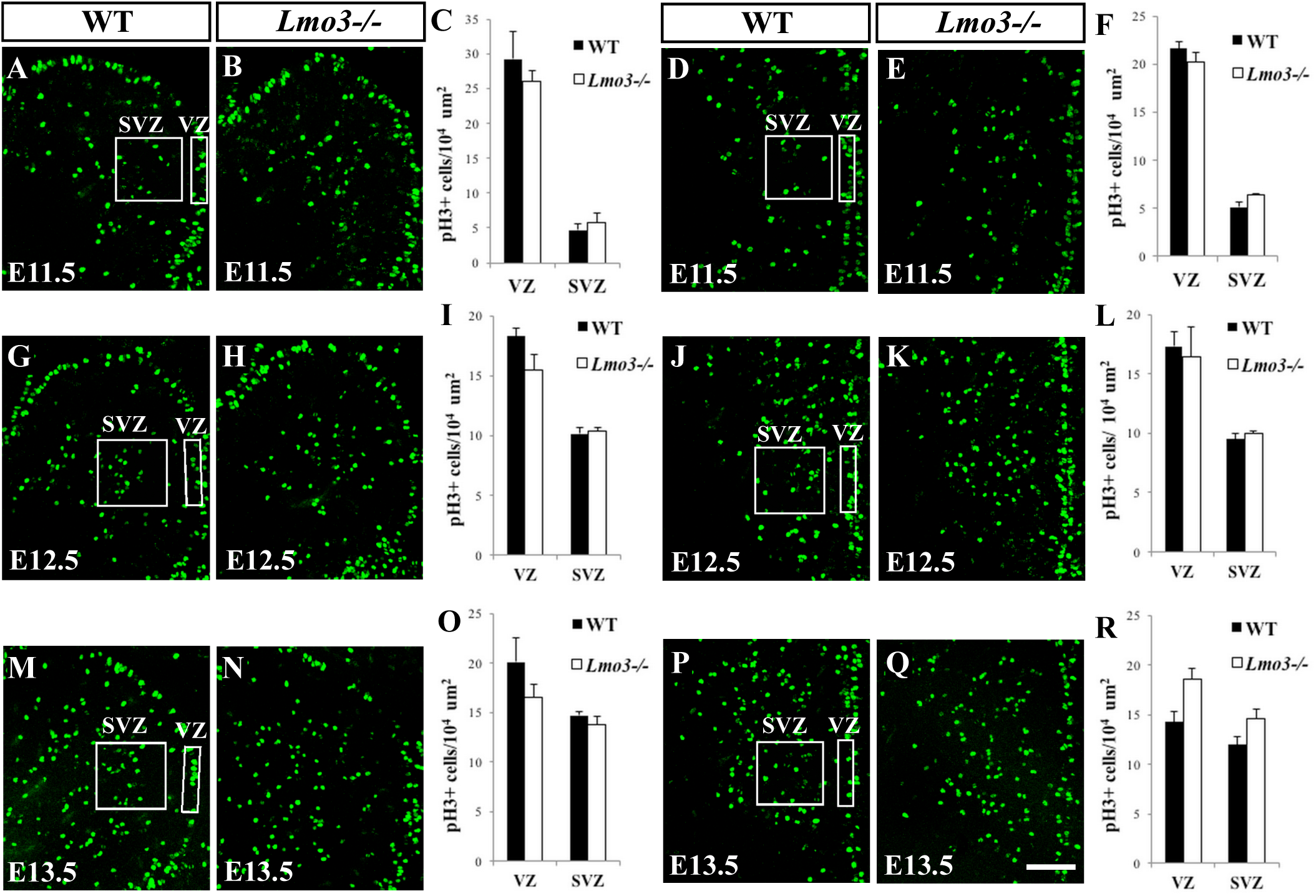
Proliferation in the WT and *Lmo3-*null MGE. Immunostaining for pH3 at E11.5 in the rostral (**A**,**B**) and caudal MGE (**D**,**E**), at E12.5 in the rostral (**G**,**H**) and caudal MGE (**J**,**K**), and at E13.5 in the rostral (**M**,**N**) and caudal MGE (**P**,**Q**) shows no significant changes in the density of proliferating cells in the VZ or SVZ in the *Lmo3-null* compared to WT (**C**, **F**, **I**, **L**, **O**, **R**). pH3^+^ cell counts were obtained from the rectangular (VZ) and square (SVZ) boxed regions in the ventral part of the MGE as indicated in these representative images, at both rostro-caudal levels and at all three time points. (*n* = between 3 and 4 for both genotypes for all graphs). [Graphs indicate mean values; error bars show SEM. Two-tailed *t* test, **C**, *p* = 0.66 for VZ and 0.597 for SVZ; **F**, *p* = 0.422 for VZ and 0.396 for SVZ; **I**, *p* = 0.156 for VZ and 0.753 for SVZ; **L**, *p* = 0.776 for VZ and 0.446 for SVZ; **O**, *p* = 0.274 for VZ and 0.4 for SVZ; **R**, *p* = 0.06 for VZ and 0.132 for SVZ]. ac, anterior commissure, MGE, medial ganglionic eminence; SVZ, subventricular zone; VZ, ventricular zone. Scale bar: 100 μm.

**Figure 7-1.**
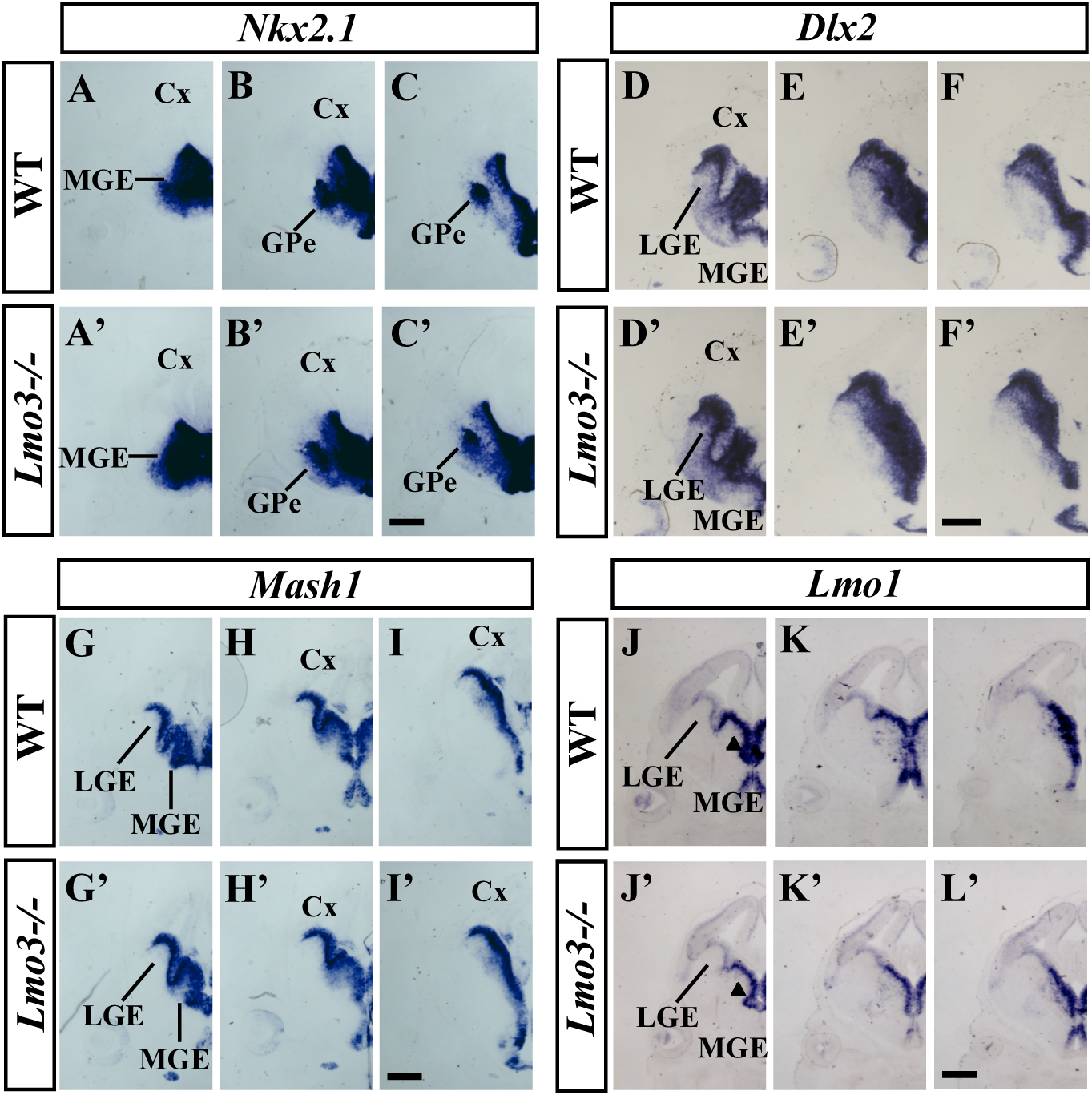
Expression of subpallial markers in the control and *Lmo3*-null mutant at E13.5. Images of three serial coronal hemisections (rostral-most on the left) show *in situ* RNA hybridization staining at E13.5 for *Nkx2.1* (**A**-**C**, **A’**-**C’**), *Dlx2* (**D**-**F, D’**-**F’**), *Mash1* (**G**-**I, G’**-**I’**) and *Lmo1* (**J**-**L, J’**-**L’**). Cx, cortex; GPe, external globus pallidus; LGE, lateral ganglionic eminence; MGE, medial ganglionic eminence. Scale bars: 500 μm.

**Figure 7-2.**
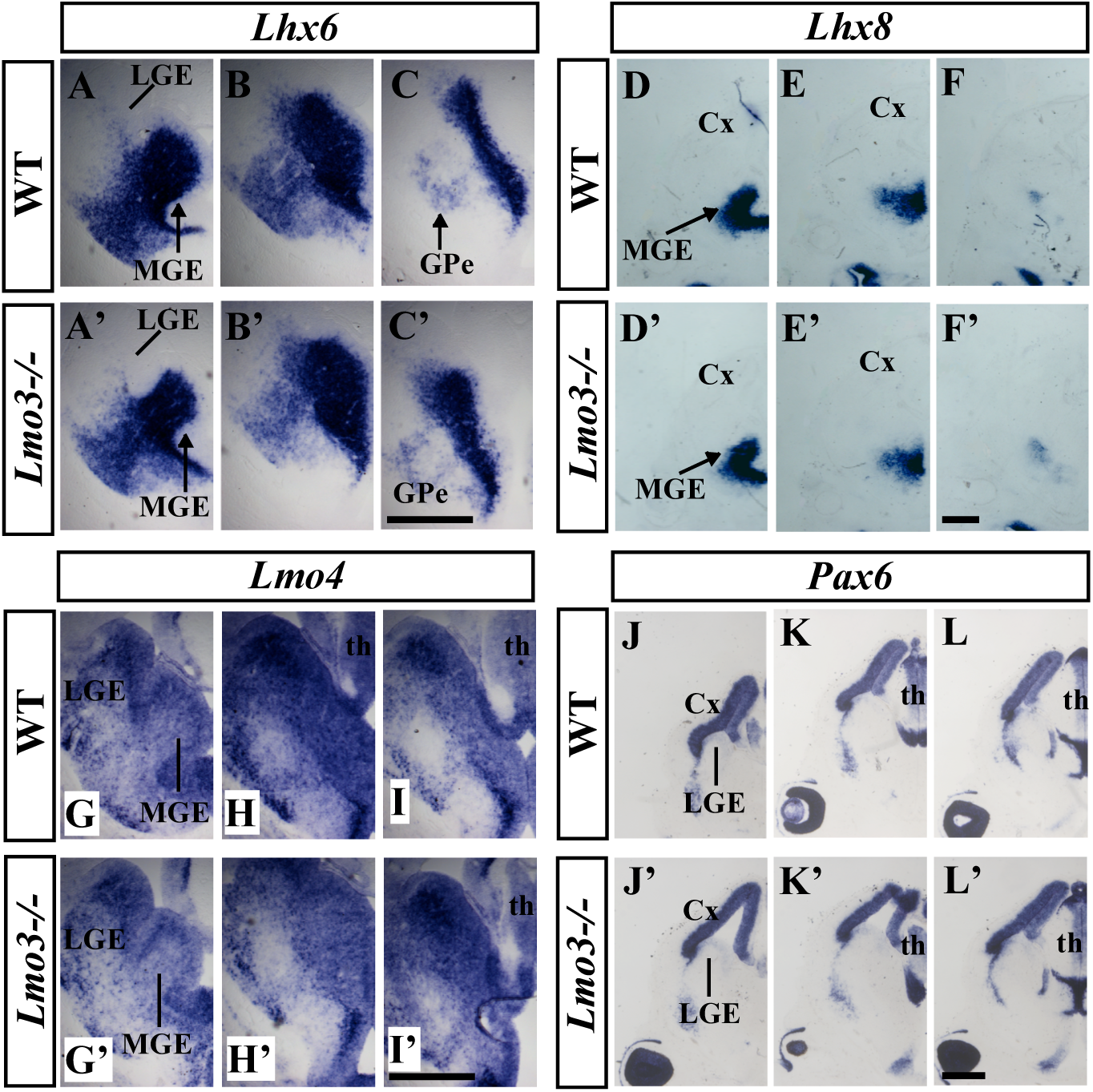
Expression of MGE and LGE markers is unchanged in the *Lmo3-*null mutant at E13.5. Images of three serial coronal hemisections (rostral-most on the left) show *in situ* RNA hybridization staining for *Lhx6* (**A**-**C**, **A’**-**C’**), *Lhx8* (**D**-**F**, **D’**-**F’**), *Lmo4* (**G**-**I**, **G’**-**I’**) and *Pax6* (**J**-**L**, **J’**-**L’**). Cx, cortex; GPe, external globus pallidus; LGE, lateral ganglionic eminence; MGE, medial ganglionic eminence; th, thalamus. Scale bars: 500 μm.

**Figure 7-3.**
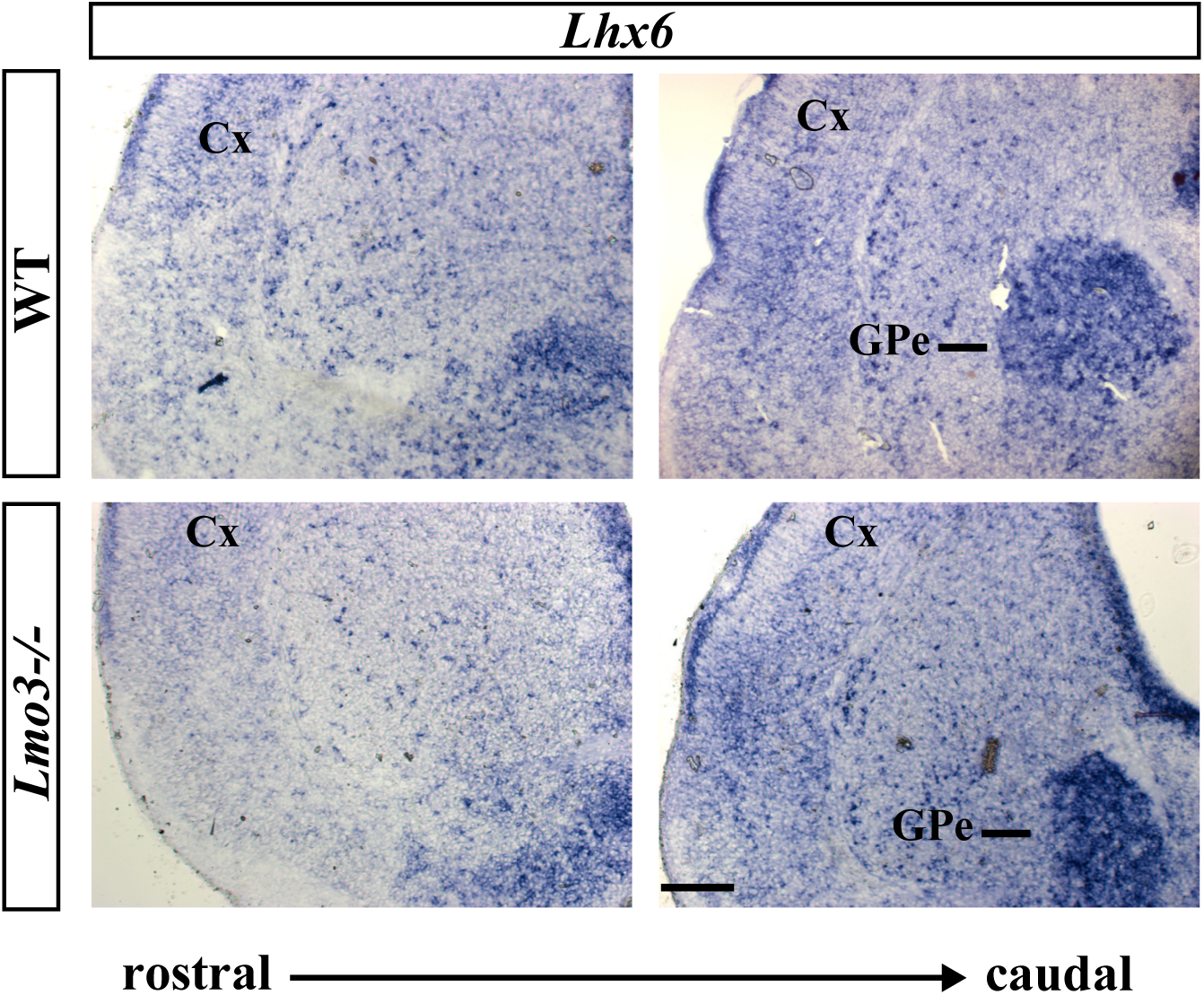
Expression of *Lhx6* in the ventral telencephalon is unchanged in the *Lmo3-*null mutant at P0. Cx, cortex; GPe, external globus pallidus. Scale bar: 500 μm.

## Notes

### Competing Interest Statement

John L.R. Rubenstein is cofounder, stockholder, and currently on the scientific board of Neurona, a company studying the potential therapeutic use of interneuron transplantation.

